# Pathway elucidation of pharmaceutical cucurbitacin IIa in *Hemsleya chinensis* and high-level production of precursor cucurbitadienol in engineered *Saccharomyces cerevisiae* and *Nicotiana benthamiana*

**DOI:** 10.1101/2022.09.28.509966

**Authors:** Geng Chen, Zhao-kuan Guo, Yan Zhao, Yan-yu Shu, Lei Qiu, Shao-feng Duan, Yuan Lin, Si-mei He, Xiao-bo Li, Xiao-Lin Feng, Gui-sheng Xiang, Yang Shi, Sheng-Chao Yang, Guang-hui Zhang, Bing Hao

## Abstract

Cucurbitacin IIa is a triterpene isolated exclusively from *Hemsleya* plants, which is non-steroidal anti-inflammatory drug that function as the main ingredient of Hemslecin capsules and Supplemental Tablets in China. In this study, the biosynthetic pathway of cucurbitacin IIa was elucidated by characterization of squalene epoxidases (HcSE1, HcSE2), cucurbitenol synthases (HcOSC6) and acyltransferases (HcAT1) in *Hemsleya chinensis*. Meanwhile, cycloartenol synthases (HcOSC1), isomultiflorenol synthases (HcOSC5) and β-amyrin synthase (HcOSC2-4) involved in sterol and triterpenes biosynthesis were functionally illustrated. The high-level production of yeast the key cucurbitacin precursor, cucurbitadienol, was constructed to produce 296.37 mg/L cucurbitadienol and 722.99 mg/L total triterpenoid which is the highest yield cucurbitadienol from known engineered microbes. Moreover, production of cucurbitenol in transient expression of tobacco was employed to achieve 94.8 mg/g dry weight (dw) cucurbitenol from leaves. In this study, the key genes involved in cucurbitacin IIa biosynthesis were identified to facilitate its medical applications via biosynthetic strategy. Meanwhile, the high-level production of cucurbitadienol chassis yeast and tobacco transient expression offered a robust and sSupplemental Table substrate for pharmaceutical cucurbitacin production and screening platform for candidate genes involved in cucurbitacin biosynthesis.

## Introduction

Cucurbitacin is a group of high-oxygen triterpenoids with cucurbitane, one of the hundreds of possible cyclic triterpenoid skeletons, as a skeleton (Xu et al., 2004; Thimmappa et al., 2014), mainly distributed in the Cucurbitaceae family such as *Citrullus lanatus, Momordica charantia, Cucumis melo, Cucumis sativus, Siraitia grosvenorii*, and the traditional Chinese herb, *Hemsleya chinensis. Hemsleya* plants are traditional Chinese herbs, the main active compounds, CuIIa and CuIIb, are used to treat digestive and respiratory inflammation. However, only a few studies have focused on the biosynthesis of CuIIa and CuIIb (Zhang et al., 2022). The complex and unique structures of such compounds exhibit specific and potent bioactivities, including anti-cancer, anti-inflammatory, anti-fertility, and anti-diabetic activities (Chen et al., 2005; Hussain et al., 2019; Ren and Kinghorn, 2019). Cucurbitacin compounds are mainly found from cucurbitaceae plants and are classified into 12 classes (A-T) based on backbone oxidation groups and other characteristics (Chen et al., 2005). *H. chinensis* contains various cucurbitacins such as cucurbitacin IIa (CuIIa) and cucurbitacin IIb (CuIIb) belonging to the cucurbitacin F class (Figure 1), and have been traditionally used for the treatment of fever, pain, and inflammation symptoms in China for thousands of years. Recently, CuIIa was found to be a novel anti-cancer agent derived from *H. chinensis* (Boykin et al., 2011; Wang et al., 2018).

**Figure 1.**
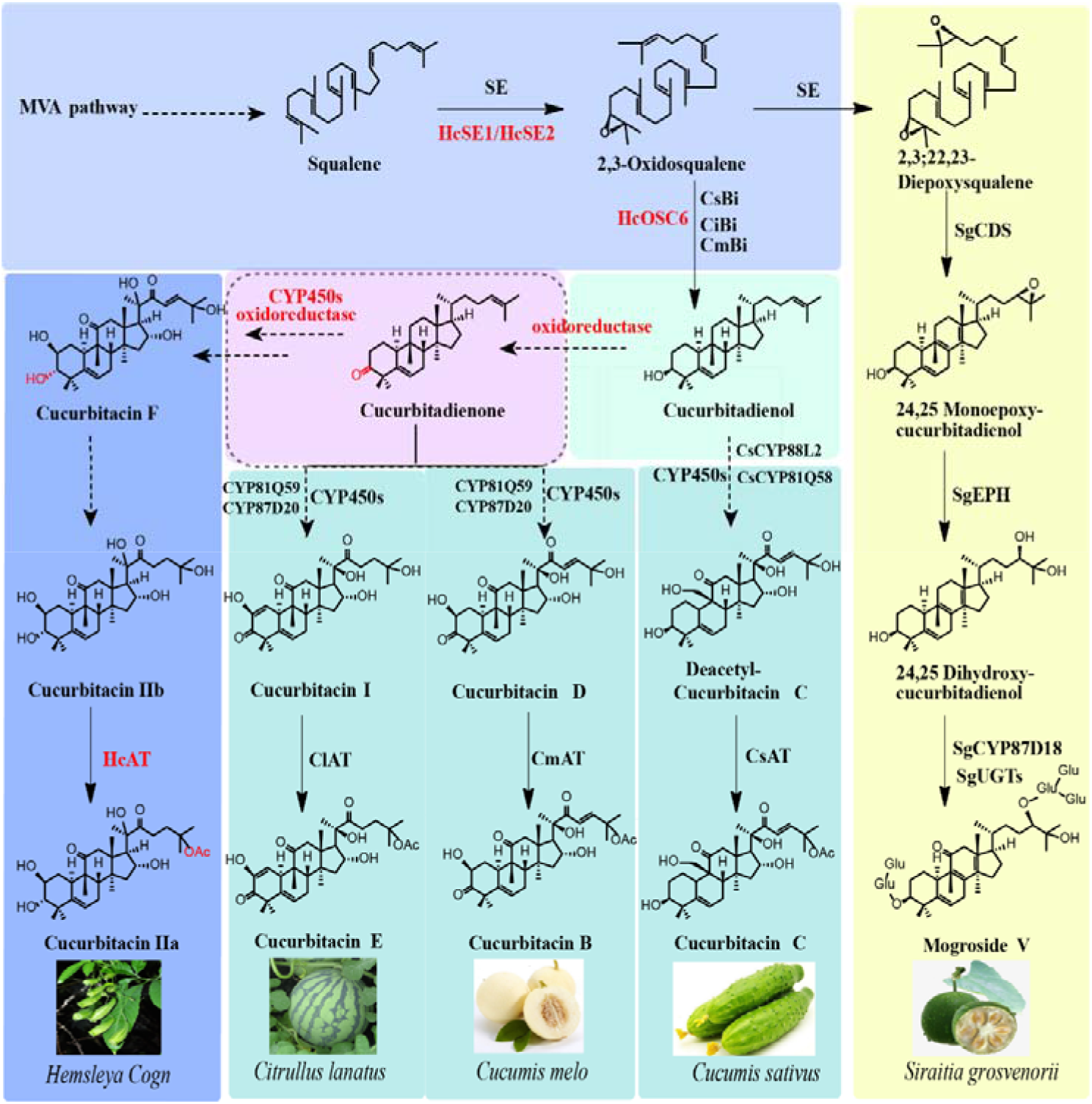
Known cucurbitacin and mogroside biosynthetic pathways and the proposed cucurbitacin F biosynthetic pathway. The red oval dotted box indicates the important reaction steps concerned with cucurbitacin F biosynthesis in this work. Each solid line arrow indicates an identified step reaction, and each dashed arrow indicates one or multiple proposed step reactions; HcSE, HcOSC, and HcAT are candidate genes from Hemsleya chinensis, which represent the biosynthetic reaction steps to be identified in this work; 3-keto-cucurbitenol in the dashed box in the pathway is proposed. Abbreviations: Bi and CDS, Cucurbitadienol synthase; EPH, epoxide hydrolase; UGT, glycosyltransferases; cyp450, cytochromes P450; SE, squalene epoxidase; OSC, oxidosqualene cyclases; AT, Acetyltransferase.

In plants, the first step in triterpene biosynthesis begins with the cyclization of 2,3 squalene or 2,3:22,23-dioxysqualene to form a triterpene backbone, and the substrate is mevalonate pathway (MVA) biosynthesis (Dewick, 2002), squalene is first catalyzed to 2,3-oxysqualene by squalene epoxidase (SE), or sequentially oxidized to 2,3:22,23-Squalene dioxide, both of which can be used as substrates for cyclization (Phillips et al., 2006). Previous results show that only three squalene epoxidases (CpSE1, CpSE2 and CpSE3) are responsible for the formation of 2,3-oxidosqualene and 2,3:22,23-dioxidosqualene in *Cucurbita pepo* (Dong et al., 2018). Cyclization is catalyzed by oxidosqualene cyclases (OSCs) that produce many triterpenes skeletons. Several *OSC* genes exist within a single plant that are responsible for the formation of characteristic triterpenes. Four *OSC* genes cuol-(McCBS), isomultiflorenol- (McIMS), β-amyrin-(McBAS) and cycloartenol-synthase (McCAS) were found to catalyzed by different triterpenes in *M. charantia* (Takase et al., 2019). Moreover, cyclization of oxidosqualene takes place in two different conformations. The chair-chair-chair conformation (CCC) that produces majority of triterpenes such as *β*-amyrin and isomultiflorenol, while the chair-boat-chair (CBC) conformation produce cycloartenol and cuol, precursors for plant sterols and cucurbitacins (Thimmappa et al., 2014), respectively (Figure 2). In *S. grosvenorii*, the cuol synthase catalyses the cyclization of 2,3-oxidosqualene and 2,3:22,23-dioxidosqualene to produce cuol and 24,25-epoxycuol (Dai et al., 2015; Itkin et al., 2016). The triterpene skeletons are further modified by cytochromes P450, acetyltransferases, and glycosyltransferases to produce various triterpenoid and saponins (Sawai and Saito, 2011). In Cucurbitaceae, biosynthesis of bitter substances (cucurbitacin) and sweet glycosides (mogroside V) are derived from 2,3-oxidosqualene which has been widely utilized. Generally, at least seven steps of oxidation and acetylation of cuol are required to obtain various cucurbitacins. Meanwhile, in *S. grosvenorii*, 2 steps of oxidation and 5 steps of glycosylation are needed to produce mogroside V (Itkin et al., 2016; Zhou et al., 2016) (Figure 1).

**Figure 2.**
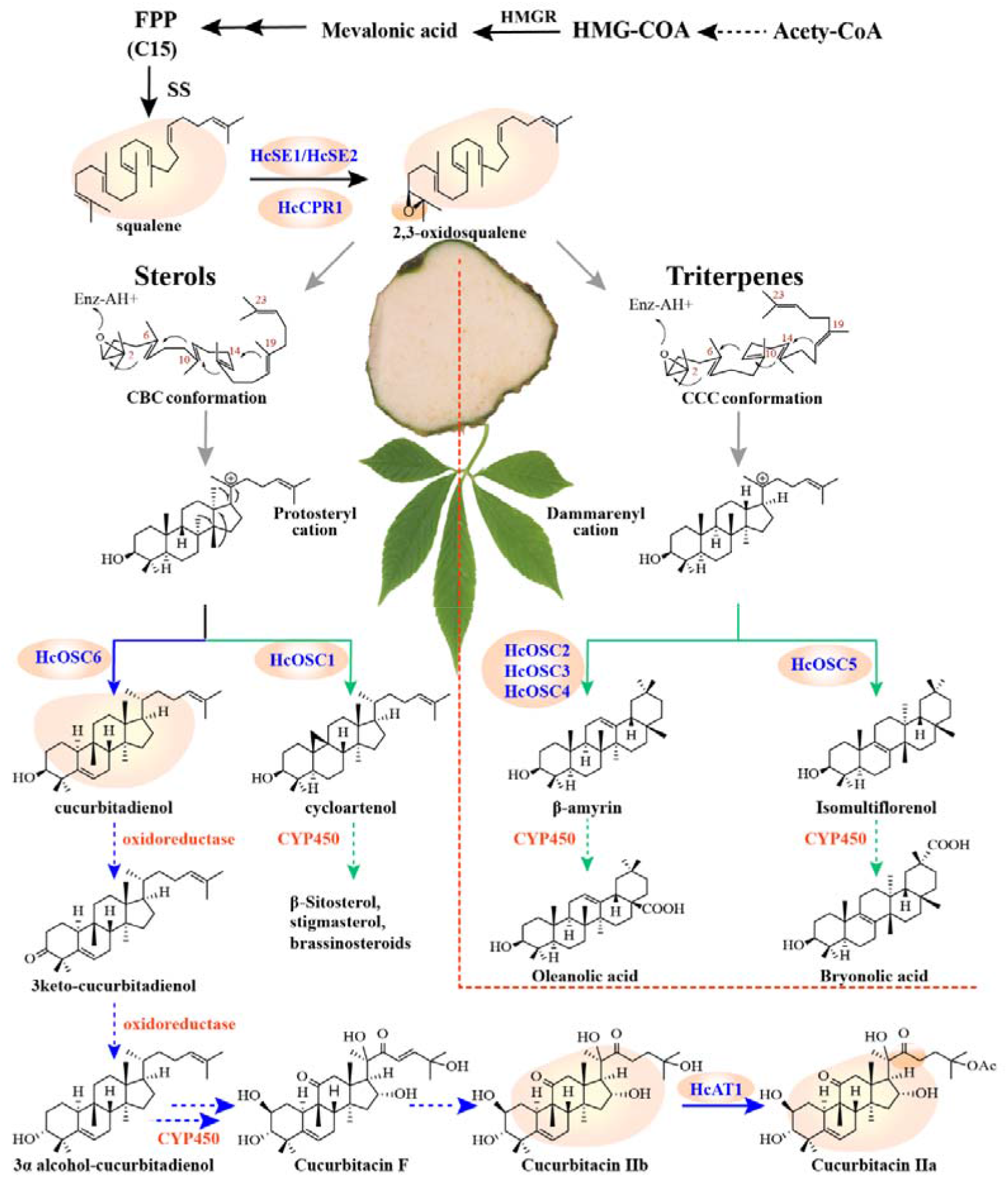
Proposed biosynthetic pathway for sterols and triterpenes in *Hemsleya chinensis*. The blue arrows indicate the proposed cucurbitacin F biosynthetic pathway, green arrows indicate proposed phytosterol and pentacyclic triterpene biosynthetic pathways; and each dotted arrow indicates one or multiple proposed reactions step; Solid arrows indicate have been identified reactions, those gene which were identified in this study are shown in blue, unidentified genes are shown in red. Enzyme abbreviations: SE, squalene epoxidase; OSC, oxidosqualene cyclases; CYP450, cytochrome P450 monooxygenases; AT, Acetyltransferase. Other abbreviations: CBC, chair-boat-chair; CCC, chair-chair-chair.

Cuol is a key starting substrate for cucurbitacin biosynthesis, further oxidized and acetylated modified multiple positions of cuol, such as C2, C7, C11, C16, C19, C20, C22, C23, C24, and C25, to produce numerous cucurbitacins, which indicates that a large amount of CYP450s are involved in these modifications (Chen et al., 2005). The biosynthetic pathways of curcurbitacins are gradually being revealed in some cucurbitaceous crops, Two *CYP450s, CsCYP88L2* and *CsCYP81Q58*, that participate in cucurbitacin C biosynthesis have been identified in *C. sativus (Shang et al., 2014*). In *C. melo* and *C. lanatus, Cm890* and *Cl890* were found to be catalysts for C11 carboxylase and C20 hydroxylase, and *Cm180* and *Cl180* for C2 hydroxylase (Zhou et al., 2016). Moreover, it was found that cucurbitacin B and cucurbitacin I was acetylated from cucurbitacin D and cucurbitacin E through the catalysis by *CmAT* and *ClAT* (Figure 1). Three new P450 genes from *M. charantia* involved in cucurbitacin biosynthesis, *McCYP81AQ19* for cuol C23 hydroxylase, *McCYP88L7* for C19 hydroxylase, C5-C19 ether-bridged products, and *McCYP88L8* for C7 hydroxylase (Takase et al., 2019).

Cucurbitacins are widely known for various biological activities; its low content in plants and complex extraction process has limited its prospects for use in the pharmaceutical market. Synthetic biology provides new strategies for these high value natural drugs. In many cases, synthesis in a heterologous host is required to produce these compounds in industrially relevant quantities. Sufficient precursor cuol is required for heterologous production of cucurbitacins and validation of pathway candidate genes in metabolic engineering. It is particularly important to construct chassis cells with high yield of cuol to increase the yield of cucurbitacins. In a previous study, the transformed yeast strain EY10 carrying the plasmid PYES2+CsCPR (Csa1G423150) + Csa3G903540 produced 10 mg/L of 19-hydroxy-cuol in shake flask fermentation (Shang et al., 2014). Increasing the flux of triterpenoid precursors has been successfully used to boost triterpene production (Wu et al., 2012; Dong et al., 2013). Modular optimization and overexpression of key genes in the relevant pathways to increase metabolic flux of precursors has been proposed (Wang et al., 2019). In a plant heterologous expression system, transient expression in *N. benthamiana* mediated by agroinfiltration is an efficient synthetic biological platform to produce plant triterpenes (Dong et al., 2013; Reed and Osbourn, 2018). Previous studies have shown that *CpSE2* and *CpCPQ* from *C. pepo* were transiently co-expressed in *N. benthamiana* with the production of cuol (0.02 ng/g dw) which is still challenging to break the cuol production limitation (Dong et al., 2018).

Therefore, in this study, we functionally characterized key *SEs*, *OSCs*, and *ATs* from *H. chinensis* for the first time, and gene-to-gene co-expression analysis with the Cytoscape software was conducted to provide clues to C-3 keto or C-3α-hydroxyl formation of cucurbitacin (Figure 2). Furthermore, we constructed a high-level cucurbitenol-producing yeast strain as the chassis cell to identify key candidate modification genes involved in cucurbitacin biosynthesis, providing the possibility to produce novel cucurbitacins or derivatives thereof through metabolic engineering.

## Results

### Isolation and sequence analysis of candidate *HcSEs, HcOSCs*, and *HcATs* based on *H. chinensis* transcriptomic analyses

Despite neither genomic nor transcriptomic data for *H. chinensis* being available, we performed transcriptional sequencing and analysis of the root, tuber, stem, leave, and flower of *H. chinensis*. We obtained a total of 50,061,733 reads and assembled them into 52,923 contigs (Supplemental Table S2). BLAST analyses were performed to annotate 32,742 of the contigs using the NCBI non-redundant, Swiss-Prot, Kyoto Encyclopedia of Genes and Genomes (KEGG), and Clusters of Orthologous Groups (COGs)/ Eukaryotic Orthologous Groups (KOGs) protein databases. Transcripts of *SEs* (*HcSE1, HcSE2* and *HcSE3*), OSCs (*HcOSC1, HcOSC2, HcOSC3, HcOSC4, HcOSC5* and *HcOSC6*), and *ATs* (*HcAT1, HcAT2, HcAT3* and *HcACT4*) candidates were screened via homology-based searches of the transcriptome. Three putative *SE* genes were cloned from the *H. chinensis* cDNA library. The *ORFs* of the three *SE* genes range from 1545–1575 nucleotides in length and separately encode 522, 524, and 514 amino-acid polypeptides with calculated molecular masses of 56.90 kDa, 57.25 kDa and 55.90 kDa, respectively. The computed isoelectric points (pI) of the predicted proteins are 8.87, 8.70, and 8.53 (Supplemental Table S3). *HcSE* protein sequences were submitted to the TMHMM Server (http://www.cbs.dtu.dk/services/TMHMM/) to predict transmembrane helices. A single transmembrane helix was predicted at the N-terminus of both the *HcSE1* and *HcSE2* protein, and four putative transmembrane helices were predicted to be distributed evenly at the N- and C-termini of the *HcSE3* protein (Figure S3; Supplemental Table S3). Functional domain analysis showed that the motif XHXHGXGXXGXXXHXXH(X)_8H_, where “X” is any residue and “H” is a hydrophobic residue, was identified in the N-terminal part of the *SE* sequences by comparing the HcSEs to the characterized *A. thaliana* and *T. wilfordii* SEs used to screen out the *H. chinensis* SEs (Figure S4). This motif is part of the Rossmann fold that binds the cofactors flavin adenine dinucleotide (FAD) and NADP^+^ in FAD^-^ and NAD(P)H-dependent oxidoreductases, respectively. The conserved motif NMRHPLTGGGMTV, known to bind squalene as a substrate in rat SE was also found in the aligned sequences (Rasbery et al., 2007) (Figure S4). Thus, the predicted proteins, corresponding to the cloned *HcSEs*, contain the typical motifs for squalene epoxidases. All six *HcOSCs* were homologous based on the multiple sequence alignments. The deduced amino acid sequences of these six *HcOSCs* showed 42.71–68.93% similarity (Figure S5). All six *HcOSCs* possessed the DCTAE motif, which is involved in substrate binding (Ito et al., 2013), and four QW motifs characteristic of the *OSC* superfamily (Figure S6). The QW motifs might be involved in stabilizing carbocation during cyclization (Srisawat et al., 2019). In addition, *HcOSC2, HcOSC3*, and *HcOSC4* have the MWCYCR motif, which is predicted to be a highly conserved motif of *β*-amyrin synthase (Zhou et al., 2019)

### Correlation of candidate *HcOSCs* and *HcATs* expression with CuIIa and CuIIb distribution in different tissues

Differential gene expression analysis and hierarchal clustering was performed using available *H. chinensis* RNA sequencing data (http://medicinalplants.ynau.edu.cn/transcriptomics/213). *HcOSC6, HcAT1*,and *HcAT2* showed the highest expression in the tuber (Figure S2A). Meanwhile, the tuber of *H. chinensis* harbored significantly more CuIIa and CuIIb than the root, flower, stem, and leaf (Figure S1A; Figure S1C).To determine whether the gene expression levels of the *HcSEs, Hc*OSCs, and *Hc*ATs in *H. chinensis* were correlated with CuIIa, CuIIb, and Oleanolic acid accumulation, the relative transcript levels of the *HcSEs*, *HcOSCs*, and *HcATs*, and the CuIIa and CuIIb content levels were determined across all sampled tissues. The relative expression levels of *HcOSC6* were highest in the tuber (Figure 3C), which also contained the highest CuIIa and CuIIb (1.43 mg/g and 0.19 mg/g dw). The relative expression levels of *HcAT1* and *HcAT2* were high in the tuber and root (Figure 3B), which were also the tissues with high CuIIa and CuIIb content.

**Figure 3.**
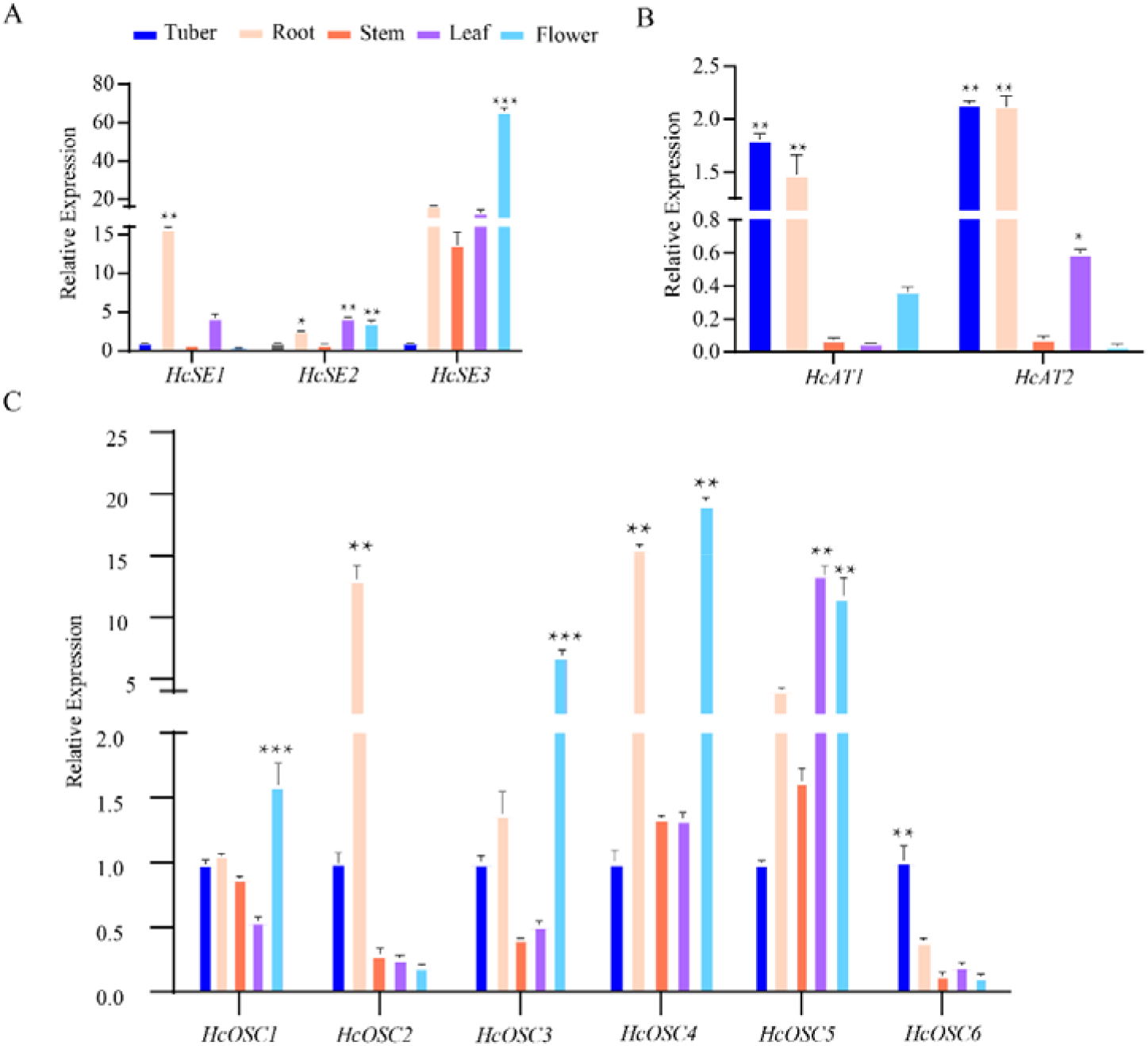
Analysis of tissue expression of *HcSEs, HcOSCs* and *HcATs* genes. A, Analysis of tissue expression of *HcSEs.* B, Analysis of tissue expression of *HcOSCs*. C, Analysis of tissue expression of *HcATs*. The gene expression was normalized by *Hc18s*. Data are presented as mean ±SD (n = 3).

### Phylogenetic tree construction of *HcSEs, HcOSCs*, and *HcATs*

The results of phylogenetic analysis showed the evolutionary position of *HcSEs* among the chosen SEs downloaded from the NCBI database (Figure S7). The *HcSEs* were clustered into a clade with other selected plant *SEs*, while other *SEs* were clustered into a mammal or fungus clades. Previous studies have reported that plant SEs can be divided into two groups, SEs and SE-like groups (Song et al., 2019). The *HcSE1–3* was classified into the SE group, which indicated that these SEs could catalyze squalene to produce 2,3-oxidosqualene. We constructed a phylogenetic tree of *HcOSCs* and characterized *OSCs* from other plant species. *Hc*OSC1 clustered within the clade of previously characterized cycloartenol synthases; *HcOSC2*, *Hc*OSC3 and *Hc*OSC4 clustered in the known characterized β-amyrin synthases; *Hc*OSC5 clustered with previously characterized isomultiflorenol *OSCs* from *Luffa aegyptiaca;* and *HcOSC6* clustered with previously characterized cuol synthases from cucurbitaceae (Figure 4). Using the sequences of BAHD-ATs identified from other species (Supplemental Table S4), we also identified 47 different putative BAHD-ATs in the transcriptome of *H. chinensis* (Irmisch et al., 2018). The results of the phylogenetic analysis showed that these 47 sequences clustered into four of the five known BAHD-AT clades, with the majority falling into clade I and V. *HcAT3* and *HcAT4* were clustered in clade I with specific AT (Csa5G639480), which was previously a negative control in cucumber leaves (Shang et al., 2014). *Hc*AT1 and *Hc*AT2 were clustered in clade III with previously characterized CsaAT2 (Csa6G088700) from *C. sativus* and CmAT1 (Melo3C022373) from *C. melo* (Figure S8).

**Figure 4.**
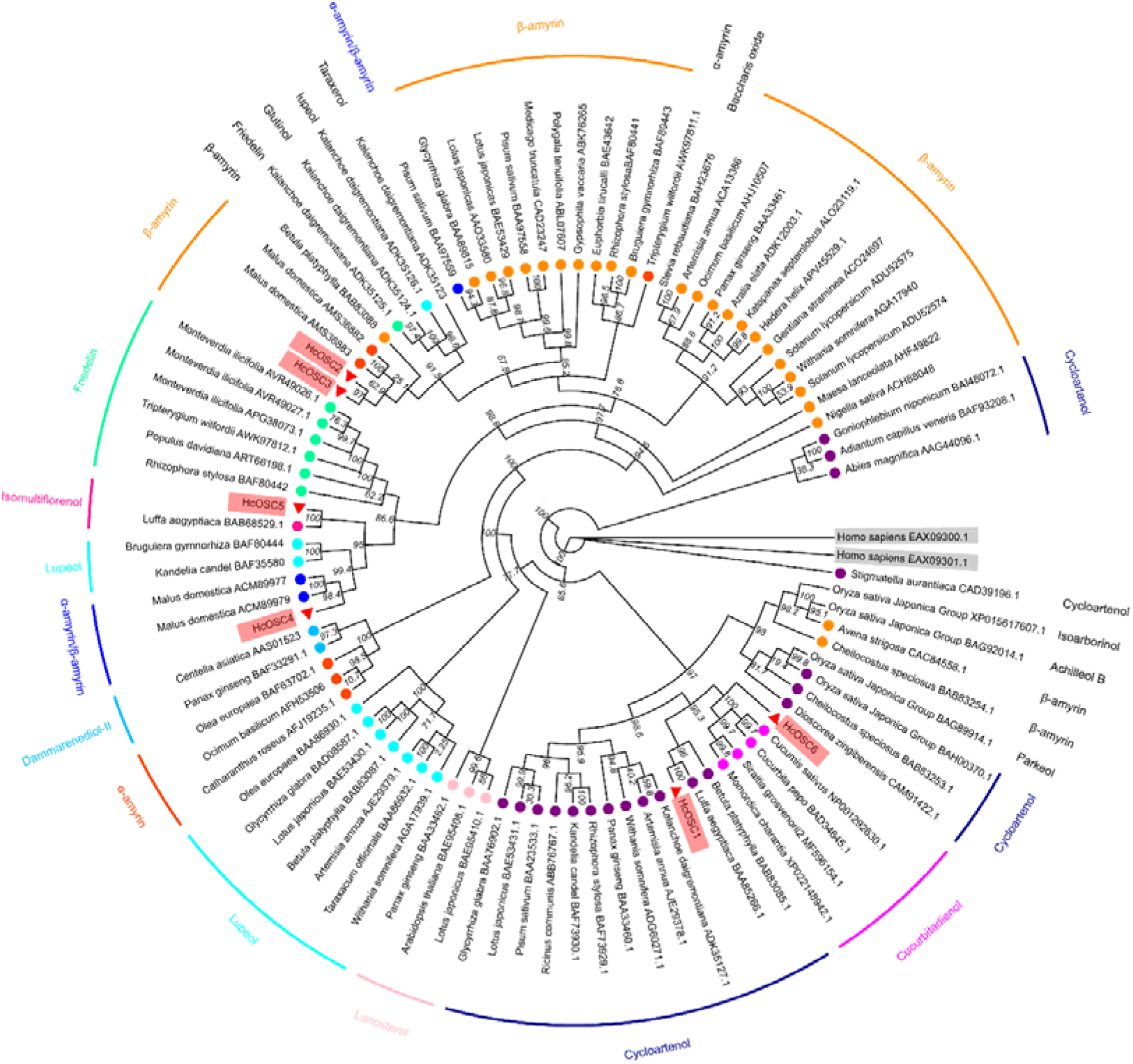
Phylogenetic analysis of oxidosqualene cyclases. Complete amino acid sequences of OSCs were obtained from the GenBank database. The maximum-likelihood (ML) phylogenies tree was built by IQ-tree software, and the number of bootstrap replications was 1000. including HcOSCs. β-Amyrin synthases, darkOrange; α-Amyrin synthases, OrangeRed; Lupeol synthases, aqua; Lanosterol synthases, Pink; Cycloartenol synthases, darkblue; Isomultiflorenol OSCs, DeepPink, Cucurbitadienol synthases, Magenta, multifunctional OSCs, blue . HcOSC1, HcOSC2, HcOSC3, HcOSC4, HcOSC5 and HcOSC6 are highlighted in gray red and red triangle marks the HcOSCs. Two human OSC gene sequences used as an outgroup are highlighted by the gray rectangle.

### Characterizations of HcSEs and HcATs

The constructed pMal-*Hc*SE1-3 plasmid was transformed into Transetta (DE3) chemically competent cells and induced at a low temperature to reduce the formation of inclusion bodies. After induction, two bands at approximately 100 kDa (56.90 kDa HcSE1 protein and 57.25 kDa HcSE2 protein plus the 42.5 kDa maltose-binding protein) were observed in the supernatant. However, no HcSE3 protein was observed in the supernatant, and protein expression was verified using analysis by SDS-PAGE (Figure S9A). To increase the solubility of the protein, 66 amino acids at the N-terminus of HcCPR1 were truncated to exclude a predicted transmembrane region (Song et al., 2019). The remaining sequence was inserted into the pET-32a vector with expression under the control of the T7-promoter. SDS-PAGE revealed that the truncated HcCPR1 was 90.65 kDa (72.65 kDa HcCPR1 with an 18-kDa fusion tag) expressed in the supernatant, protein expression was verified using analysis of SDS-PAGE and protein gel blot (Figure S9B). HcSE1 and HcSE2 function was determined by GC-MS analysis of the products from the incubation of purified recombinant HcSE1/HcSE2 and HcCPR1 proteins with squalene and NADPH(Song et al., 2019). The GC results revealed that a new peak appeared at 16.02 min (Figure 5A), which was identified as 2,3-oxidosqualene according to the characteristics of the primary ion peaks (Figure 5B). However, as a control experiment, when the amount of exogenous squalene was increased 2,3-oxidosqualene was not detected in the extracts without addition of the HcCPR1.

**Figure 5.**
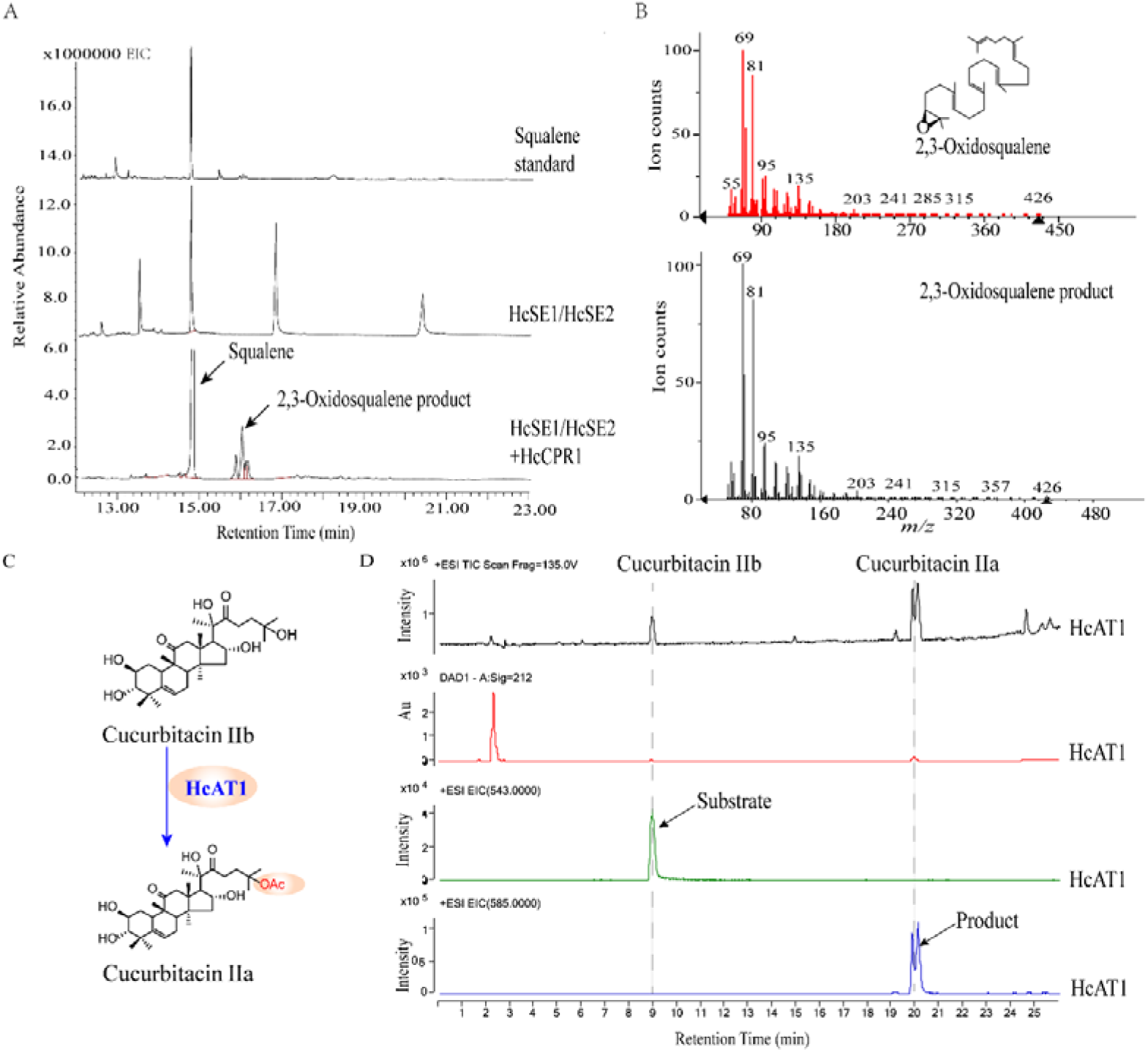
Enzyme activity of recombinant HcSE, HcCPR1 and HcATs. (A) GC results of the HcSE1, HcSE2 and HcCPR1 enzymatic reaction. (B) MS results revealed the primary ion peak of the 2,3-oxidosqualene standard and the enzymatic product. (C) Catalyzes the final step of cucurbitacin IIa synthesis. (D) LC-MS results of the HcATs enzymatic reaction.

The HcATs sequence was inserted into the pET-32a vector with expression under the control of the T7-promoter which expressed the target protein. All four HcATs were expressed in *E. coli* and protein expression was verified by analysis of SDS-PAGE and protein gel blot (Figure S9C). Two bands at approximately 68 kDa (47.81 kDa, *Hc*AT1 protein, 47.55 kDa *Hc*AT2 protein, 52.38 kDa, all with the 18-kDa fusion tag protein) (Figure S9C). According to the deduced route, the protein encoded by HcAT1 may be involved in catalyzing the acetylation of the hydroxyl group at the C-25 position (Figure 5C). CuIIb was used as the substrate for *in vitro* enzymatic reaction, and HPLC detection showed that HcAT1 could catalyze acetyl groups onto CuIIb to produce CuIIa, However, *Hc*AT2, HcAT3, and HcAT4 protein failed to demonstrate the catalytic activities. Therefore, *Hc*AT1 successfully catalyzed the final step of CuIIa (Figure S10A). To further confirm that the product produced by the *in vitro* enzymatic reaction is CuIIa, the product and the standard CuIIa were detected by LC-MS. The peak time of the reaction product and the standard product CuIIa was at 20.02 min, and the results of the characteristic peaks are consistent, further confirming that the generated product is CuIIa (Figure 5D, Figure S10B).

### Functional characterization of HcOSCs in yeast

To identify the functions of the putative OSCs from *H. chinensis*, the complete ORFs of all the HcOSCs were sub-cloned into the vector pYES2 and then transformed into lanosterol synthase-deficient yeast (GIL77). In parallel, the empty pYES2 vector was transformed into the yeast mutant to serve as a negative control. After culturing and induction, the metabolite extracts were analyzed by GC-MS and NMR spectra to confirm the products. Based on the results of GC and NMR spectra analysis showed that the GIL77 strain expressing HcOSC1 produced an evident product at 22.10 min, which exhibited identical retention time and mass spectral characteristics to those of the purified standard cycloartenol (Figure S11, Figure S13). GC-MS analysis showed that the extracts from yeast harboring HcOSC2, HcOSC3, and HcOSC4 contained β-amyrin, their products all exhibited identical retention time at 24.80 min and mass spectral characteristics to those of the authentic standard of β-amyrin (Figure 6C and 6D). The GIL77 strain expressing HcOSC5 produced an evident product at 24.20 min, of which product exhibited identical retention time and mass spectral characteristics to those of the purified standard isomultiflorenol (Figure S12). Fortunately, the GIL77 strain expressing HcOSC6 produced an evident product by analysis of TLC (Figure S17A), of which product exhibited identical retention time at 22.60 min and mass spectral characteristics to those of the purified standard cuol (Figure 6A and 6B). Meanwhile HcOSC1, HcOSC5, and HcOSC6 products were analyzed by NMR spectra (Figure S13, S14, and S15; Supplemental Table S5), their NMR data are consistent with previous literature reports (Yoshida et al., 1989; DavidáNes, 1991; Isaev, 1995). These result from GC-MS and NMR spectra analysis proved that HcOSC1 was cycloartenol synthases, HcOSC5 was isomultiflorenol synthases, HcOSC6 was cuol synthases, and HcOSC2-4 were a β-amyrin synthase.

**Figure 6.**
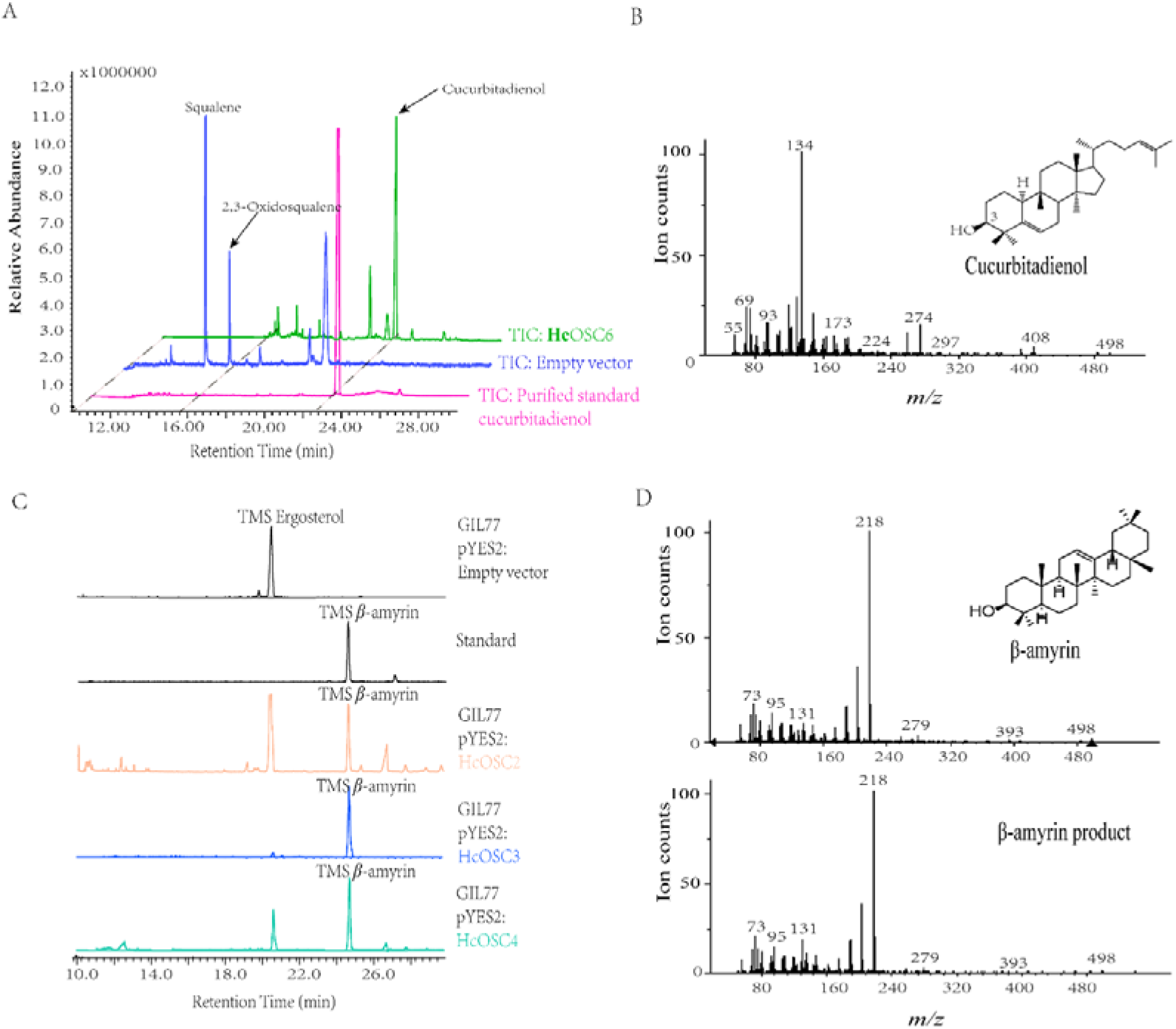
Gas chromatography-mass spectrometry (GC-MS) analysis of the products in yeast strains harboring the *Hc* OSCs (*Hc*OSC2,3,4 and *Hc*OSC6) from *Hemsleya chinensis*. A, The gas chromatography identification of cucurbitadienol produced by HcOSC6 compared with the standard cucurbitadienol obtained by preparative liquid chromatography. B, The mass spectra of the HcOSC6 product peak and purified standard cucurbitadienol. C, The gas chromatography identification of cucurbitadienol produced by HcOSC2,3,4 compared with the standard β-amyrin. D, The mass spectra of the HcOSC2,3,4 product peak and standard β-amyrin.

### Construction of cuol producing chassis yeast

Cuol is the key precursor of cucurbitane-type triterpenoids (Figure 1), the accumulation of cuol directly determine the yield of cucurbitacin in cell factory. Ten enzymes are required to synthesize substrate 2,3-oxidosqualene from acetyl coenzyme A (acetyl-CoA) in the MVA pathway. After formation of 2,3-oxidosqualene, *HcOSC6* subsequently catalyze cuol production. Therefore, we aimed to overexpress these 12 genes to increase the yield of cuol in chassis yeast (Figure 7A). The 12 genes were divided into two modules, the designated upstream-module included seven upstream genes (*ERG10, ERG13, ERG12, ERG8, ERG19, IDI*, and *tHMG1*) that converted acetyl-CoA to the isoprenoid building blocks isopentenyl phosphate (IPP) and dimethylallyl phosphate (DMAPP). The designated downstream module, contained the remaining five downstream genes (*ERG1*, *ERG20*, *ERG9, synHcCPR1* and *synHcOSC6*), which converted IPP and DMAPP into the target product cuol. *HcOSC6* and *HcCPR1* were codon-optimized for expression in yeast and were renamed *synHcOSC6* and *synHcCPR1*, respectively. HMG-CoA reductase catalyzes the rate-limiting step in the MVA pathway, multi-copy overexpression of tHMG1 could increase the production of different terpenoids (Wang et al., 2019). Therefore, we added an additional copy of tHMG1 to the second group (Figure 7A). After integrating the downstream-module into the delta DNA site of the yeast strain BY4742, the resulting strain named CUOL01-1, produced 133.21 mg/L cuol and 173.02 mg/L ergosterol from glucose in shake flasks (Figure 7B and 8C; Supplemental Table S7). This demonstrated that our design is feasible, because only a single-step integration was required to increase the cuol yield by > 10-fold compared with that of the GIL77 cell strain expressing HcOSC6 (Figure 7C). In the next step, the upstream-module was integrated into the Yprc-delta15 site of CUOL01-1, the resulting strain yielding strain CUOL02-2. Its cuol titer further increased to 296.37 mg/L in shake flasks (Figure 7C; Supplemental Table S11). The cuol titer of strain CUOL02-2 was 55%greater than that of strain CUOL01-1 (Figure 7C and 7D), which indicated that introducing the upstream module boosted the metabolic flux toward the MVA pathway to promote the accumulation of cuol.

**Figure 7.**
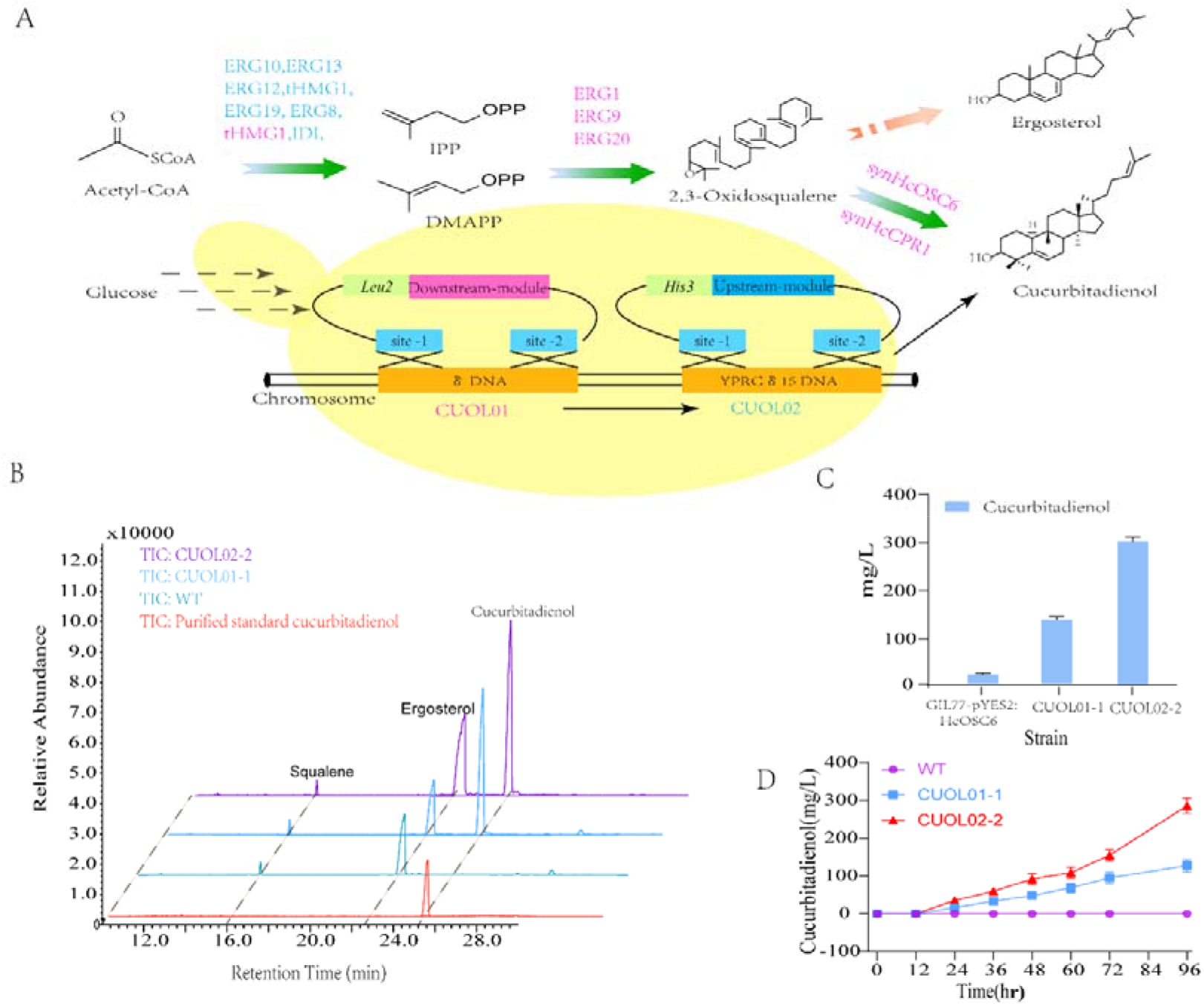
Schematic biosynthesis pathway of cucurbitadienol in engineered yeast. Blue-colored genes form the upstream-module, Magenta-colored genes form the downstream-module, green arrow for cucurbitadienol synthesis pathway. IPP isopentenyl pyrophosphate, DMAPP dimethylallyl pyrophosphate, Cuol01 chassis strain 1.0, Cuol 02 chassis strain 2.0. B, GC-ms analysis of squalene, Ergosterol and cucurbitadienol (CUOL) produced by WT(control), CUOL01-1, and CUOL02-2. C, Quantification of cucurbitadienol(CUOL) produced by Cuol 01-1, and CUOL02-2 and GIL77-pYES2. The error bars indicate the SEMs of three biological replicates. D, Quantification of cucurbitadienol(Cuol) produced by WT(control), Cuol 01-1, and Cuol 02-2 in different time gradients(96 hour), The error bars indicate the SEMs of three biological replicates.

**Figure 8.**
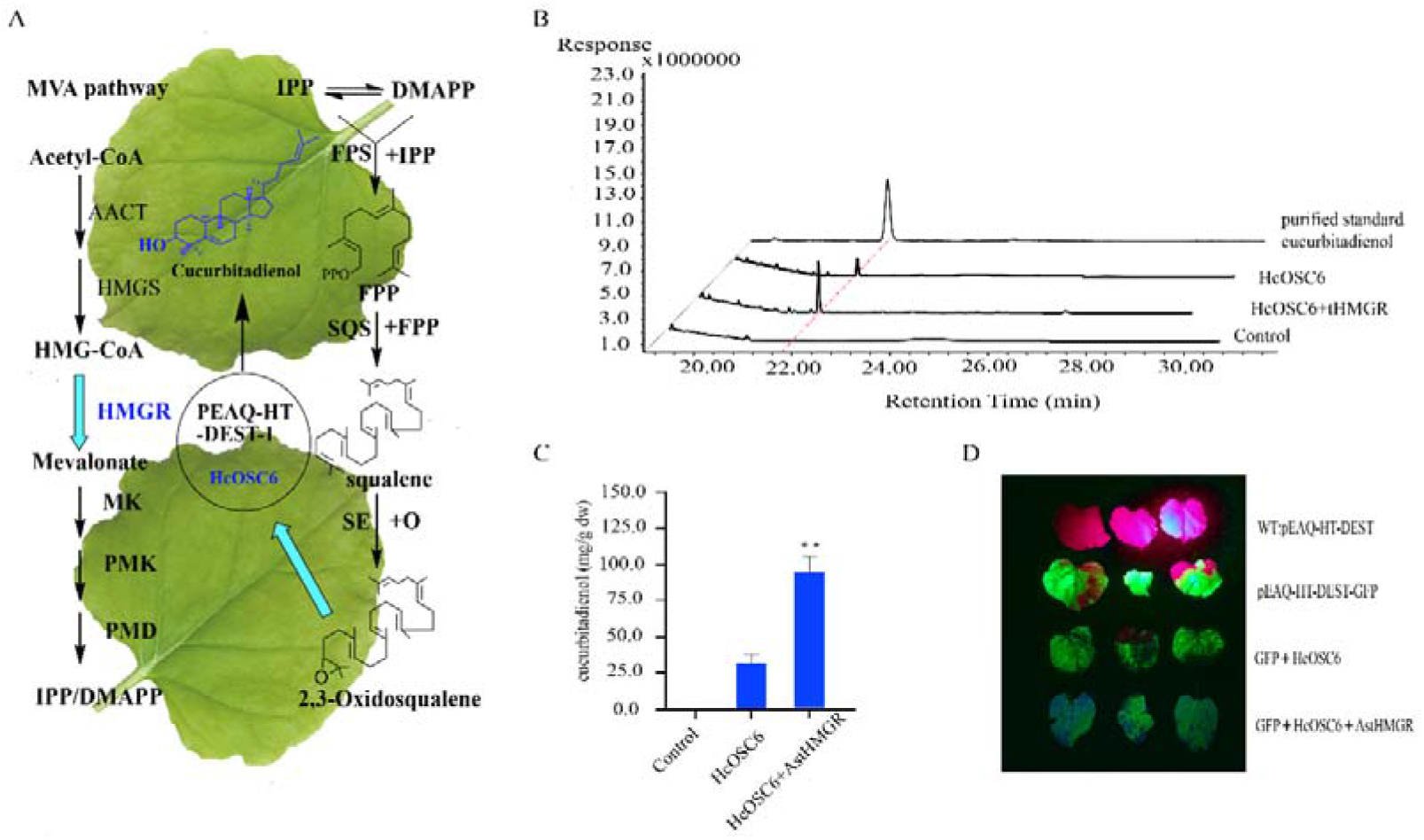
Co-expression with AstHMGR gives enhanced intermediates cucurbitadienol levels in *N. benthamiana*. A, Biosynthesis of triterpenes cucurbitadienol occurs via the mevalonate pathway. B, Total ion chromatograms (TICs) for extracts from leaves expressing HcOSC6 with either AstHMGR. C, cucurbitadienol content of tobacco leaves co-expressing HcOSC6(cucurbitadienol synthase) with AstHMGR (mean, three biological replicates ± se; control, empty vector (EV) or pEAQ-HT-DEST with GFP gene; *, P< 0.05; n.s., not significant.

### Efficient production of cuol in *N. benthamiana* leaves

Regarding the yeast expression system, HcOSC6 appears to produce cuol specifically. The *N. benthamiana* transient expression system was used to further evaluate the function of HcOSC6 and establish the plant chassis to produce cuol. Cuol content of tobacco leaves co-expressing *HcOSC6* cuol synthases with green fluoresce protein (GFP), AstHMGR1 through improvement of the agroinfiltration process (Reed et al., 2017), pEAQ-HT-DEST1-GFP were used as background controls. After three days of transient expression in *N. benthamiana*, the GFP fluorescent protein can be easily observed using a LUYOR-3415RG hand-held lamp (Luyor Corporation, Shanghai, China) to visually identify transient osmotically expressed gene events. We observed a single major peak at 20.02 min in the gas chromatogram of *N. benthamiana* leaves transiently co-expressing *HcOSC6* cuol synthases with *GFP*, *AstHMGR1*, and the same peak was not observed in the control (Figure 8B). The peak was identified as cuol by comparison of its mass fragmentation pattern with that of the purified standard cuol (Figure 8). Since precursor availability may limited, we co-expressed *HcOSC6* with upstream mevalonate pathway genes *AstHMGR1* to determine the effects on cuol production. The total content of cuol was determined to be 28.32 mg/g(dw) when only *HcOSC6* was expressed, and when *HcOSC6* and *AstHMGR1* were co-infiltrated in tobacco the total content was 94.8 mg/g(dw), demonstrating the efficient production of cuol in *N. benthamiana*.

### Gene co-expression analysis of oxidatively modified genes involved in CuIIa biosynthesis

The expression patterns of all genes were obtained from transcriptome sequencing with WGCNA analysis, which were divided into 17 modules based on the similarity of expression patterns (Figure S18A and 18B). Gene Ontology (GO) and KEGG annotation analysis of the genes obtained revealed the terpenoid backbone biosynthesis pathway (ko00900), sesquiterpenoid and triterpenoid biosynthesis pathway (ko00909) and steroid biosynthesis pathway (ko00100) pathway were enriched. Set in grey60, coral1, brow4, and other modules (Figure S18C and 18D). With functionally charactered SEs, OSCs,and ATs in this study, candidate genes involved in oxidative modification of the cucurbitacin biosynthesis pathway were identified from these terpenoid synthesis modules (Supplemental Table S6). By calculating the correlation between genes and the connectivity in the WGCNA, the core genes identified as significantly related to *HcSEs, HcOSCs* and *HcATs* were displayed in the Cytoscape software. The result indicated these core genes may be involved in cucurbitacin biosynthesis (Figure S2B). Notably, the oxidative-modifying enzymes involved in cucurbitacin biosynthesis are clustered with the characterized *HcOSC6* and *HcATs* with a strong correlation. Moreover, there is a significant correlation between Oxidoreductase1 and *HcOSC6* and *HcAT1* (cor>0.95 and p<.001) (Supplemental Table S8).The candidate oxidative modification genes were excavated including: 6 CYP450 genes, 3 Oxidoreductases and 3 alcohol reductases, named as *CYP81Q59, CYP81Q58, CYP87D19, CYP87D20, CYP90B1, CYP720B1* and *Oxidoreductase1*, *Oxidoreductase2*, *Oxidoreductase3* and *CAD-1*, *CAD-2*, *CAD-3*, respectively (Figure S2B; Supplemental Table S6). The heat map results showed that these cucurbitacin biosynthesis pathway modifying enzymes were associated with *HcOSC6* and *HcATs* (Figure S2A).

## Discussion

Cucurbitacins have been widely utilized in the past decades due to their various biological activities. Elucidating the relevant enzymes for cucurbitacin metabolism in plants provides the possibility to produce cucurbitacins via a synthetic strategy. The key precursor, cuol plays an essential role in cucurbitacin biosynthesis in both plants and heterogeneous expression systems. The content of cuol directly affects the yield of cucurbitacin production in engineering yeast and transient tobacco. Here, *HcSE1*, *HcSE2*,and *HcOSC6* involved in cuol biosynthesis were characterized from *H. chinensis* which requires a two-step catalyzation from squalene to cuol. HcSE1 and HcSE2 were successfully expressed and purified in the prokaryotic system. However, the ORF sequence of the *HcSE3* is approximately 30bp less than HcSE1 and HcSE2, and four putative transmembrane helices were predicted to be distributed evenly at the N- and C-termini of the HcSE3 protein (Figure S10; Supplemental Table S3). In addition, the helices of HcSE1 and HcSE2 were predicted to start from the inside of the endoplasmic reticulum (ER) membrane (ER lumen), while the helices of HcSE3 were predicted to start from the outside of the ER membrane (cytosol) with 6.9% of total at the N-terminus (Supplemental Table S3). Overall, these results suggest that HcSE proteins might have different membrane topologies that may result in HcSE3 protein expression failure.

Functional identification of 6 HcOSCs in lanosterol synthase-deficient yeast revealed different product specificity. HcOSC1, a cycloartenol synthase showed an evolutionary conserved essential function in the plant sterol pathway (Corey et al., 1993). HcOSC2/3/4 acted as a monofunctional OSC which could cyclize 2,3-oxidosqualene to β-amyrin (Figure 6C and D). Surprisingly, HcOSC5 was found to be an isomultiflorenol OSC that converted 2,3-oxidoqualene to isomultiflorenol as an intermediate of pentacyclic triterpene, which subsequently converted into bryonolic acid(Hayashi et al., 2001). Bryonolic acid was shown to accumulate exclusively in the roots and cultured cells of various cucurbitaceous plants (Cho et al., 1992). This is the first report of isomultiflorenol synthase as a potential regulatory enzyme controlling the biosynthesis of the bryonolic acid skeleton in *H. chinensis;*bryonolic acid has not been previously reported in the *Hemsleya* genus. These findings indicated that HcOSC2/3/4/5 were responsible for the catalysis of pentacyclic triterpenes, while the product of HcOSC1 produced cycloartenol and phytosterols, such as campesterol and sitosterol, which are biosynthesized via cycloartenol and catalyzed by cycloartenol synthase (CAS) in higher plants (Ohyama et al., 2009) (Figure 2). HcOSC6 is a key enzyme for cucurbitacin biosynthesis, catalyzing 2,3-oxidosqualene to form cuol which is the starting substrate for CuIIa and other cucurbitacins. In a previous study, 4 OSCs were reported as cuol (*Mc*CBS), isomultiflorenol (*Mc*IMS), β-amyrin (*Mc*BAS), and cycloartenol synthase (*Mc*CAS) in *M. charantia* (Takase et al., 2019).

The robust chassis strain for sufficient precursor supply is an important strategy for increasing the yield of natural products via a synthetic biological approach. It is necessary to boost metabolic flux of the key substrate and optimize the rate-limited enzymes in the pathway. In this study, to construct a high yield cucurbitenol-producing yeast strain, the biosynthetic flux of cucurbitenol (Cuol) was systemically enhanced via over expressing all the 10 pathway genes. The pathway was divided into two modules and modular engineering strategies were adopted for chassis strain construction. A single-step integration of the downstream-module resulted in the production of more than 130 mg/L of cuol after 96 h in shake flasks (Figure 7) which indicated the feasibility of modular engineering. The subsequent integration of the upstream module resulted in 296.37 mg/L cuol production and total triterpenoid production reached 722.99 mg/L (Figure 7). Cuol synthase (CSB) from *C. sativus* (*Cs*Bi) have been utilized to produce cuol in yeast. The integration of HMGR, ERG20, ERG9, and ERG1 into the chromosome of the yeast strain and their over expression could increase the yield of cuol by approximately 10-fold, calculated from relative average peak area with yeast plasmid system. Here, we constructed the cuol-producing chassis strain CUOL02-2, which could produce 296.37 mg/L cuol in a shake flask system. To date, this is the highest cuol yield in engineered microbes. However, during the fermentation process, it was found that ergosterol has a high accumulation in the native pathway (up to 426.62 mg/L). Thus, improvement of the synthesis capacity of the downstream pathway will continuously increase product yield. For instance, increasing cuol synthase activity by screening for more natural sources or preforming directed evolution via protein engineering. Moreover, the cuol synthase could be placed under a higher strength promoter, and optimization of fermentation conditions to further increase the yield and conversion of cuol in yeast.

The tobacco heterologous production platform was performed to produce cuol in this study. Currently, Colchicine, saffron crocins Momilactone, and taxadiene-5α-ol are heterologously synthesized in tobacco (Reed et al., 2017; Li et al., 2019; Martí et al., 2020; Nett et al., 2020; De La Peña and Sattely, 2021; Nett and Sattely, 2021). Transient expression of Agrobacterium infiltration into *N. benthamiana* provides a new platform for efficient and rapid preparation of monomeric drugs. Here, the improvement of the agroinfiltration process vacuum infiltration was used to express AstHMGR and HcOSC6. To increase our capacity for effective leaf infiltration we referred to a previous report and performed efficient vacuum infiltration of 6–8 plants simultaneously (Reed et al., 2017). Preliminary experiments using a reporter construct showed that this method resulted in clear GFP expression in infiltrated leaves (Figure 8D). The infiltration of *N. benthamiana* plants with *A. tumefaciens* strains carrying CPMV-HT expression was constructed for the *HcOSC6* to produce 28.32 mg/g(dw) of cuol. Previous reports showed co-expression of SAD and AstHMGR1 produced 4-fold higher amounts of β-amyrin than when SAD was singly expressed (Reed et al., 2017). Further co-overexpression of upstream mevalonate pathway genes AstHMGR1 and *HcOSC6* resulted in the triple production of 94.8 mg/g(dw) cuol (9.48% dw of the leaves). Therefore, co-expression of HcOSC6 and AstHMGR was employed to significantly increase the yield of cuol. In this study, we provide a high yield of cuol yeast chassis and tobacco transient expression system which could be used for efficient cucurbitacin production and screening platform for candidate genes involved in cucurbitacin biosynthesis.

Culla is the main pharmacological component in *H. chinensis*; however, its biosynthesis pathway is still not clear. Our results showed that HcAT1 could catalyze acetyl groups onto CuIIb to produce CuIIa, which indicates that *HcATs* have the same function as *Cm*AT, *Cs*AT, and *Cl*AT in cucurbitacin B, C, and E biosynthesis (Figure 5 and S10). To identify modification genes involved in CuIIa biosynthesis, the expression profiling datasets from five different tissues of *H. chinensis* obtained from RNA sequencing analysis were used. We particularly focused on CYP450s and oxidoreductases, that catalyze oxygenation reactions and oxidoreduction of triterpenoid skeleton. The result revealed OSC and AT genes that showed characteristic tissue-dependent expression patterns (Figure 3 and S2A). *Hc*OSC6 particularly encoding cuol synthase, showed the highest expression in tubers rather than in other tissues.

CYP450s and oxidoreductases involved in CuIIa biosynthesis were expected to demonstrate a similar expression pattern. Thirty-five genes in the triterpene biosynthesis pathway were annotated with a BLASTX search. Thirty-two genes that showed a highly similar expression pattern with the *HcOSC6* and *HcAT1* were selected using the Cytoscape software (ver3.7.1), to construct the correlation networks. The result showed 12 modified genes involved in cucurbitacin biosynthesis with *Hc*OSC6, *HcAT1*, and *HcAT2* that have been characterized are strong correlated including six P450 genes (CYP81Q59, CYP81Q58, CYP87D19, CYP87D20 CYP90B1, CYP720B1), three dehydrogenases (*Oxidoreductase1*, *Oxidoreductase2*, *Oxidoreductase3*), and three cinnamoyl-CoA reductases (CAD-1, CAD-2, CAD-3) (Figure S2D; Supplemental Table S2). In the further research, the function of the 12 modified genes involved in CuIIa will be verified and applied into our efficient cucurbitacin production platform to elucidate the complete biosynthesis pathway of CuIIa and achieve high-yield CuIIa production.

## Conclusions

We characterized two squalene epoxidases (HcSE1 and HcSE2), oxidosqualene cyclases HcOSC6 and one acyltransferase (HcACT1) involved in CuIIa biosynthetic pathways in *H. chinensis*. The functional analysis revealed that both HcSE1 and HcSE2 could catalyze oxidize squalene to form 2,3-oxidosqualene, following cyclization by HcOSC6 to from cucurbitadienol which is the key precursor for cucurbitacin biosynthesis. Cucurbitacin was subsequently modified by P450s and oxidoreductases before the last catalyzation by HcACT1 to produce CuIIa. The function of cycloartenol synthase HcOSC1, 3 β-amyrin synthases (HcOSC2-4), and isomultiflorenol synthase HcOSC5 involved in sterol and triterpenes biosynthesis in *H. chinensis* were also illustrated in this work. Moreover, the key precursor of cucurbitacins, cuol, its biosynthetic pathways was constructed in engineered yeast to produce 296.37 mg/L cuol and 722.99 mg/L total triterpenoid. To date, this result is the highest cuol yield from known engineered microbes. Meanwhile, we achieved production of cuol in transient expression of tobacco leaves with the yield of 94.8 mg/g(dw).

## Materials and Methods

### Plant materials

Mature (6-year-old) *H. chinensis* plant tissues were collected from the Kunming in Yunnan Province, China. Five tissue samples of root, tuber, stem, leaves, and flowers were harvested from plants, and stored in liquid nitrogen until use.

### UPLC analysis of CuIIa, CuIIb and Oleanolic acid content in plant tissues

Samples of 5 tissues (tubers, roots, leaves, stems and flowers) collected from *H. chinensis* were vacuum freeze-dried (Christ ALPHA 1-2, Germany) and ground to a fine powder. About 0.5 g of each tissue sample was taken, 3 copies each. Accurately weighed, placed in a stoppered conical flask, accurately added 25 mL of 70% methanol, weighed, ultrasonically treated (power 180 W, frequency 40 kHz) for 40 minutes, left to cool, weighed again, and used 70 % methanol to make up the lost weight and shake well. Extracted three times with ethyl acetate, combined and concentrated. Dissolve with methanol, dilute to 10 mL, take an appropriate amount of the supernatant, filter it through a 0.22 μm microporous membrane, and take the filtrate for ultra-high performance liquid chromatography (UPLC) (Agilent) analysis.

Content analysis was performed using an Agilent 1290 Infinity II High Performance UHPLC System. The chromatographic column was a Phenomenex Kinetex C18 analytical column (4.6×100 mm, 2.6 μm), and the temperature was maintained at 30 °C. The mobile phase for the detection of CuIIa and CuIIb is 0.2% phosphoric acid aqueous solution (A)-acetonitrile (B), gradient elution: 0 ~ 15 min, 25% ~ 33%B; 15 ~ 20min, 33% ~ 40%B; 20 ~ 24 min, 40% ~ 60% B; 24 ~ 28 min, 60% ~ 90% B; detection wavelength 212 nm. The mobile phase for the determination of oleanolic acid was 0.2% phosphoric acid aqueous solution (A)-acetonitrile (B), gradient elution: 0 ~ 10 min, 40% ~ 69% B; 10 ~ 15 min, 69% ~ 69% B; 15 ~ 20 min, 69%~80%B; detection wavelength 201 nm. The flow rate was 0.8 mL·min-1, and the injection volume was 5 μL.

### Total RNA isolation, cDNA synthesis, library construction, and sequencing

Total RNA was isolated from H. chinensis plant tissues using HiPure HP Plant Total RNA Kit (Magen, China). The extracted RNA was used to synthesize cDNA according to the kit instructions. The kit used for qPCR experiments was PrimeScript™ II 1st Strand cDNA Synthesis Kit (TaKaRa, Japan), HiScriptII 1st Strand cDNA Synthesis Kit (Vazyme Biotech Co., Ltd.) was used for gene cloning experiments. Sequencing was performed using an Illumina HiSeq TM 4000 from Gene Denovo Biotechnology Co. (Guangzhou, China). After purifying the cDNA fragments, the ends were repaired, poly(A) was added, and ligated to Illumina sequencing adapters. Pre-sequencing assessment of RNA quality was checked by analysis on an Agilent 2100 Bioanalyzer (Agilent Technologies, Palo Alto, CA, USA).

### Cloning of *HcSEs, HcOSCs*, and *HcATs* coding sequences

*SEs*, *OSCs*, and *ATs* candidates were identified from the *H. chinensis* transcriptome database by BLAST searching for known *SEs*, *OSCs* and *ATs* from cucurbitaceae and other plants. The protein-coding gene sequences of candidate genes were obtained from the transcriptome data of *H. chinensis*and primers were designed for the amplification of gene sequences of *HcSE*, *HcOSC* and *HcAT* (Supplementary Table S1). Coding sequences were obtained according to the manufacturer’s instructions for the Phanta Max Super-Fidelity DNA Polymerase Kit (Vazyme Biotech Co., Ltd.), and coding sequences were recovered by gel and ligated into expression vectors according to the manufacturer’s instructions for the ClonExpress II One One Step Cloning Kit (Vazyme Biotech Co., Ltd.). Ltd.), the gene sequences were recovered by gel and ligated into the expression vector according to the manufacturer’s instructions. Finally, the gene sequences were cloned into the vector and transferred to E. coli DH5α by thermal excitation for positive sequencing and sequence correctness.

### Sequence and phylogenetic analyses

In order to obtain the open reading frames (ORFs) and amino acid sequences of HcSEs, HcOSCs and HcATs and to align and identify protein functional domains, the online tool ORF Finder (http://www.ncbi.nlm.nih.gov/gorf) was used in this study. /gorf.html), DNAMAN software was used for multiple sequence alignment, and Interpro (www.ebi.ac.uk/Tools/InterProScan) was used to identify functional domains. The amino acid sequences of OSCs of other species were downloaded from the National Center for Biotechnology Information (NCBI) database and aligned by ClustalW. The phylogenetic tree is a maximum-likelihood (ML) phylogenetic tree constructed by the bootstrap method using IQ-tree software with 1000 replicates. (Nguyen et al., 2015)

### Tissue expression patterns and quantitative real-time PCR

Total RNA from 5 tissues of H. chinensis was reverse transcribed for quantitative real-time PCR (qRT-PCR) according to the manufacturer’s procedures and principles for the HiScript III RT SuperMix for qPCR (+gDNAwiper) Kit (Vazyme Biotech Co., Ltd.). First stop cDNA. Specific primers for qRT-PCR (Supplementary Table S1) were designed using online software (http://www.primer3plus.com/) and gene expression was detected on the Applied Biosystems QuantStudio 5 real-time PCR system (Applied Biosystems, New York, USA). PCR conditions were as follows: initial incubation at 95°C for 3 minutes, followed by 45 cycles of 95°C for 3 seconds and 60°C for 30 seconds, followed by melting curve cycling. 18SrRNA was used as a reference gene for relative quantification. Relative transcript abundance was evaluated using the 2^-ΔΔCt^ method (Livak and Schmittgen, 2001) with measurements performed in triplicate from three biological replicates.

### Functional identification in *E. coli*

The entire coding region of HcSEs was cloned into the maltoprotein-tagged pMal-c2x vector by homologous recombination using the ClonExpress II One Step Cloning Kit(Vazyme Biotech Co.,Ltd.) according to the instructions . The N-terminal transmembrane domain of HcCPR1 was truncated at 66 amino acid residues(M1-V66) was constructed on pET-32a vectors using the same method as above. The vector recombination reaction solution was transferred to competent cells Transetta (DE3) (Shanghai Weidi Biotechnology Co., Ltd.), coated on LB medium plate, the temperature was maintained at 37 °C, and cultured overnight, and then the bacteria water PCR sequencing confirmed the positive, Select positive single colonies, inoculate into LB liquid medium containing 100 mg/L ampicillin for overnight culture, transfer the culture to 400 mL LB medium containing 100 mg/L ampicillin, and inoculate at 37 °C with Shake at 180 rpm until the OD600 (optical density) reaches 0.6. Expression of HcSEs and HcCPR1 was induced with isopropyl β-D-1-thiogalactopyranoside (IPTG) at a concentration of 0.1 mM by shaking at 160 rpm at 18 °C for 16 h.

Protein purification and enzymatic detection of HcSE and HcCPR1 were performed as previously described(Song et al., 2019). HcSE was purified by MBPTrap HP affinity column (Abbkine, China) and eluted with 10 mM maltose, HcCPR1 was purified by Ni-NTA agarose affinity column and 80 mM imidazole, all buffer systems were Tris-HCl. Recombinant proteins were detected by 10% (w/v) sodium dodecyl sulfate polyacrylamide gel electrophoresis (SDS-PAGE) by 10% SDS-PAGE (w/v) gel electrophoresis, and BSA was used as quantification Standard determination of protein concentration. The enzymatic activity of HcSEs is determined by adding HcCPR1 to the reaction system to provide hydrogen peroxide to provide oxygen to catalyze the substrate squalene to produce 2,3-squalene oxide or 2,3:22,23-dioxidosqualene (Laden et al., 2000). The total volume of the enzyme reaction mixture was 1 mL, containing 40 mM squalene substrate (Sigma-Aldrich, USA), 50 ug recombinant HcSE protein, 50 μg recombinant HcCPR1 protein, 1 mM FAD, 1 mM NADPH, Triton X-100 (1%) and 50 mM Tris-HCl (pH 7.5). A negative control was used without 50 μg of recombinant HcCPR1. Extracts from the enzyme reaction mixture after overnight incubation at 25°C were analyzed by gas chromatography-mass spectrometry (GC-MS).

The protein expression, purification and enzymatic activity detection of HcATs were as described previously(Zhou et al., 2016). HcATs were successfully constructed on the pET32a vector. Positive colonies were cultured in LB medium, and cells were harvested after inducing protein expression with 0.1 mM IPTG at 16 °C for 18 hours. After disrupting the cells, the recombinant His-tagged HcAT was purified by Ni affinity chromatography in a buffer system (50 mM sodium phosphate, pH 8.0, 500 mM NaCl, 80 mM imidazole). Purified proteins were assayed by SDS-PAGE using BSA as a quantification standard. The enzyme activity of HcATs was determined by high performance liquid chromatography (HPLC) and liquid chromatography-mass spectrometry (LC-MS) to analyze whether the product CuIIa in the enzyme reaction solution, the enzyme reaction solution contained 40 μg purified HcATs, 400 μM substrate CuIIb And add buffer containing 40 mM acetyl-CoA and 50 mM sodium phosphate (pH 7.5). A negative control was used without recombinant HcCPR1 (Zhou et al., 2016).

### Immunoblotting

For detection of purified proteins, the reference method performed immunoblot detection of the proteins of HcCPR1 and HcATs(Xue et al., 2018). Proteins were prepared in loading buffer with sodium dodecyl sulfate (SDS), and then loaded on LK202 Omni-PAGE 10–12% Bis-Tris gels (EpiZyme, China) for electrophoresis. The proteins were transferred from the gel to the membrane by a high-quality wet protein transfer machine (genscript, eBlot-L1, China) with parameters set to 17 min at 50 V. Membranes were blocked in PBST-buffered saline containing 0.05% (v/v) Tween 20 and again at room temperature with mouse monoclonal His-tag monoclonal antibody (GenStar, China) containing 5% w/v skim milk After incubation in PBST buffer for 1 hour, followed by washing, followed by incubation with IgG-HRP) conjugated secondary antibody (GenStar) and washing thoroughly. Visualization was performed by StarSignal Plus Chemiluminescence Detection Kit (GenStar) (Tanon5200, China).

### Functional identification in yeast and metabolite extraction

The cloned HcOSCs were ligated to the yeast expression vector pYES2 by ClonExpress II One Step Cloning Kit (Vazyme Biotech Co., Ltd.), and the plasmids were extracted after identifying positive colonies by sequencing. The expression plasmid was transformed into yeast by electroporation, with pYES2 without HcOSC as a negative control strain. The transformation method was as described previously. GIL77 yeast cells were grown in shaking culture in YPD medium supplemented with ergosterol (20 μg/mL), heme (13 μg/mL), and Tween 80 (5 mg/mL) (Kushiro et al., 1998). Yeast cells were harvested by centrifugation. After washing treatment with water, 1 M sorbitol, and 0.1 M lithium acetate, dissolved in 1 M sorbitol, 100 μL of cells were transferred to a 0.2 cm electroporation cuvette, and 1 μg of plasmid DNA was added. Electroporation was performed using a GenePulser Electroporation System (BioRad) with the following settings: 1.5 kV, 600 Ω, 25 μF. After allowing electroporated cells to recover in YPD medium for 1 hour, transformants were screened for 2-3 days on solid medium without uracil (SC-Ura), and positive yeast strains were identified by sequencing (Dong et al., 2018). Culture of positive transformants and detection of HcOSCs products were performed as described (Zhou et al., 2019). Yeast cell culture was first shaken in SC-Ura medium containing 2% glucose for 2 days, then switched to SC-Ura medium containing 2% galactose for 1 day, and finally in 0.1 M potassium phosphate buffer (Incubate overnight in pH 7.0 with 2% glucose). Finally, the extracted products from yeast cells were analyzed by GC-MS. The extracted products could be obtained by ultrasonic extraction with methanol or by refluxing with 10 mL of 20% KOH and 50% EtOH for 5 min, and then the supernatant was extracted three times with petroleum ether.

### Purification and structural elucidation of HcOSC1, HcOSC5, and HcOSC6 products

10 liters of HcOSCs-expressing GIL77 cells were grown and pelleted, yielding 48.25 g of material. The extracted HcOSCs were obtained by saponification cleavage as previously described (Zhou et al., 2019). The extract was dissolved and separated by silica gel column chromatography (zcx. II, particle size 200-300; Haiyang, Qingdao, China) with hexane:ethyl acetate (15:1, v/v). Compounds were eluted by the x system and the combined target products were analyzed by thin layer chromatography (TLC). Further purification of the target product was performed using a reverse-phase HPLC semi-preparative column (Agilent Zorbax SB C18, 250 × 9.4 mm, 5 μm) to collect the target compound manually. In this study, Cycloartenol (20 mg), isomultiflorenol (15 mg) and cuol (55 mg) were purified from 10 L of S. cerevisiae culture expressing HcOSC1, HcOSC5 and HcOSC6, respectively. The purified compounds were analyzed by nuclear magnetic resonance (NMR).

### Construction of yeast strain producing cuol

As previously described, yeast engineering strains are generated by yeast homologous recombination (Wang et al., 2015; Wang et al., 2019). The specific step is to generate each gene expression cassette in the reconstituted pathway by PCR amplification, and the homologous sequences shared by its gene expression cassettes are used for recombinant or fusion PCR(Figure S16A; Supplemental Table S1). Fusion fragments obtained by fusion PCR are performed using adjacent genes. The fusion fragments were co-transformed into yeast strains by the lithium acetate method. Integration into the chromosomes of homologous sequences is achieved by sharing 40-75 bp of homologous sequences, and finally through the corresponding nutritional screening to generate yeast transformants.

### Yeast cultivation and metabolites extraction

YPD medium was used for yeast growth culture as in previous methods, and transformants were grown in SC solid culture corresponding to the auxotrophic marker for selection (Wang et al., 2019). Then, the positive yeast transformants were shaken and cultured, and the seed liquid was obtained and then inoculated in YPD medium for 96 hours. The fermentation broth was extracted with a 1:1 mixture of methanol and acetone and used to analyze the titer of cuol.

### Transient expression in *N. benthamiana* leaves

The cloned gene sequences of *HcOSC6* and *AstHMGR1* were subcloned into the transient expression vector pEAQ-HT-DEST1 binary vector by ClonExpress II One Step Cloning Kit (Vazyme Biotech Co., Ltd.), and then transformed into Agrobacterium tumefaciens strain The positive strains were identified by sequencing in LBA4404 (Shanghai Weidi Biotechnology). According to the engineering design (Fig. 8A), AstHMGR (HMG-CoA reductase from *A. strigosa*, GenBank accession number tHMGR – KY284573) was previously reported to promote triterpenoid production (Reed et al., 2017). As previously described methods, we co-expressed HcOSC6 with AstHMGR1 and co-introduced different strain combinations to test gene combinations (Reed et al., 2017). Briefly, after the transformed A. tumefaciens were grown in LB medium at 28°C for 24 hours, the cells were harvested with 10 mM MES (2-(N-morpholino)ethanesulfonic acid)), 10 mM MgCl2 and 100 mM acetosyringone (4′-hydroxy-3′,5′-dimethoxyacetophenone) were suspended in) to adjust the OD600 to 0.8, and then incubated in the dark with gentle shaking for 2h. Agrobacterium tumefaciens containing AstHMGR1 and HcOSC6 were infiltrated into N. benthamiana leaves at 5 weeks by a modified vacuum infiltration method, and the leaves of the plants were harvested 5 days later for triterpenoid analysis (Reed et al., 2017).

### Co-expression network analysis

The co-expression networks of 15 transcriptome data of *H. chinensis* were used were constructed using weighted gene co-expression network analysis (WGCNA) (v1.47) package in R (version 3.2.2), after filtering genes, a total of 19271 genes were used for WGCNA, gene expression values were imported into WGCNA to construct co-expression. These modules were obtained using the automatic network build function block with default settings. Using Cytoscape_3.3.0 to visualize the network to analyze gene co-expression relationships (Otasek et al., 2019).

### TLC and GC-MS analysis of extracts

For TLC analysis of target triterpenoids, yeast extract or tobacco extract was dissolved in 1 ml of ethyl acetate, the sample was loaded on TLC silica gel plate (Qingdao Haixiang, China) with a capillary tube, and the developing solution was prepared with n-hexane: Ethyl acetate or petroleum ether ethyl acetate 15:1~10:1 is developed, dyed with 10% sulfuric acid-ethanol solution and heated to observe the color development of cucurbitadienol and ergosterol.

Resuspend the extracted samples in 200 μL of extraction solvent and dry 50 μL aliquots under nitrogen. Dried aliquots were then derivatized with 50 μL of 1-trimethylsilyl (Sigma-Aldrich) and transferred to glass inserts in glass autosampler vials. GC-MS analysis was performed using a 7890B GC (Agilent) as previously described (Hodgson et al., 2019). Briefly, a 1 μL sample (inlet 250 °C) was injected in pulsed no-spigot mode (pulse pressure 30 psi), with a program consisting of oven temperature at 170 °C for 2 min, ramping to 300 °C at 20 °C/min C and 300 °C for 11.5 min. Detection was performed in scan mode (60–800 mass units) with a solvent delay of 8 min set to 7.2. Data and target compound analysis were performed using MassHunter Workstation (Agilent) software.

### HPLC and HPLC/ESIMS analysis

The analytical method used to detect the target product in the HcATs reaction is to detect the content of CuIIa and CuIIb in the plant sample, as described above. In addition, compound analysis was performed using an Agilent 1290 UPLC/6540 Q-TOF system (Agilent Technologies) equipped with an Agilent Poroshell 120 EC C18 analytical column (4.6 × 100 mm, 2.7 μm). The positive ion mode of the ion source was used, and the voltage was set to 3500 V; fragmentation voltage: 135 V; cone voltage: 60 V; RF voltage: 750 V, scanning range: 100–1000 m/z.

## Author Contributions

BH, GZ, and SCY conceived the study. GC, ZG, YS, LQ, SD, YL, SH, XB,and XF performed the experiments. YS conducted a photo shoot of plants. GC, ZG, BH and GZ the designed experiments. GC, ZG, YZ and GX analyzed the data. GC, BH, and GZ drafted the manuscript. BH, GZ, and SCY reviewed and edited the manuscript. All authors contributed to the article and approved the submitted version.

## Conflict of interest

The authors declare no competing interests.

## Accession numbers

The transcriptome reported in this paper has been deposited in Medicinal Plants multi-Omics Database(MPOD), http://medicinalplants.ynau.edu.cn/transcriptomics/213, and listed in Supplemental Table S10. The raw transcriptome sequencing data reported in this paper have been deposited in the National Center for Biotechnology Information (NCBI) database under project number PRJNA879990.

## Supplemental data

The following materials are available in the online version of this article.

Supplemental Figure S1. UHPLC analysis of content of cucurbitacin IIa, cucurbitacin IIb and oleanolic acid in *H.chinensis*.

Supplemental Figure S2. Expression patterns of HcOSC6 and other co-expressed in *H.chinensis*, as well as, gene-to-gene co-expression analysis.

Supplemental Figure S3. HcSE protein sequences were submitted to the TMHMM Server (http://www.cbs.dtu.dk/services/TMHMM/) to predict transmembrane helices.

Supplemental Figure S4. Deduced amino-acid sequence alignment of three SEs from *H.chinensis*.

Supplemental Figure S5. Amino acid sequence identity: comparison among six OSCs cloned from *H.chinensis*.

Supplemental Figure S6. Deduced amino-acid sequence alignment of six OSCs from *H.chinensis*.

Supplemental Figure S7. Phylogenetic tree of selected SEs from different species.

Supplemental Figure S8. Phylogeny of BAHD–ATs identified in transcriptome of *H.chinensis*.

Supplemental Figure S9. SDS–PAGE of the HcSEs, HcCPR1 and HcATs.

Supplemental Figure S10. HPLC and MS analysis of HcAT1 enzyme catalyzed products.

Supplemental Figure S11. GC-MS analysis of the products in yeast strains containing the HcOSC1 expression plasmids and empty vector.

Supplemental Figure S12 GC-MS analysis of the products in yeast strains containing the HcOSC5 expression plasmids and empty vector.

Supplemental Figure S13. NMR spectra of HcOSC1 product.

Supplemental Figure S14. NMR spectra of HcOSC5 product.

Supplemental Figure S15. NMR spectra of HcOSC6 product.

Supplemental Figure S16. Schematic diagram of the upstream and downstream modules of the fenugreek biosynthetic pathway and the fusion fragment of the gene expression cassette.

Supplemental Figure S17. TLC analysis of cucurbitadienol production by HcOSC6.

Supplemental Figure S18. WGCNA of differentially expressed genes.

Supplemental Table S1. Primers used in this study.

Supplemental Table S2. Summary of RNA sequencing analysis of five tissue of H.chinensis.

Supplemental Table S3. Transmembrane region prediction of HcSEs by TMHMM.

Supplemental Table S4. NCBI identification numbers of BAHD-AT sequences used in the phylogenetic analysis.

Supplemental Table S5.13C & 1H δ assignments for cucurbitadienol (HcOSC6 Product), cycloartenol (HcOSC1 Product) and (HcOSC5 Product).

Supplemental Table S6. Genes from RNA-seq analysis of H.chinensis highly correlated with HcOSC6 and HcAT1 in Cytoscape software analysis.

Supplemental Table S7. cucurbitadienol production by different strains in shake flasks.

Supplemental Table S8. Correlation analysis of oxidoreductase 1 with HcOSC6 and HcAT1.

Supplemental Table S9. Yeast strains used in this study

Supplemental Table S10. Sequence information of candidate genes and synthetic genes used in this study.

## Funding

This work was supported by Major Science and Technology Projects in Yunnan Province (2019ZF011-1), Fundamental Research Project of Yunnan(202101AS070037), Science and Technology Innovation team of Yunnan (202105AE160011), Yunnan Characteristic Plant Extraction Laboratory (2022YKZY001), the First Projects of Science and Technology Plan in the Biomedical field in 2021 (202102AA310048), and National Natural Science Foundation of China (Grant No. 81960691, 82160727).

## Parsed Citations

**Boykin C, Zhang G, Chen YH, Zhang RW, Fan XE, Yang WM, Lu Q (2011) Cucurbitacin IIa: a novel class of anti-cancer drug inducing non-reversible actin aggregation and inhibiting survivin independent of JAK2/STAT3 phosphorylation. Br J Cancer 104: 781-789**

<u>Google Scholar: Author Only Title Only Author and Title</u>

**Chen JC, Chiu MH, Nie RL, Cordell GA, Qiu SX (2005) Cucurbitacins and cucurbitane glycosides: structures and biological activities. Nat Prod Rep 22: 386-399**

<u>Google Scholar: Author Only Title Only Author and Title</u>

**Cho HJ, Tanaka S, Fukui H, Tabata M (1992) Formation of bryonolic acid in cucurbitaceous plants and their cell cultures.**

**Phytochemistry 31: 3893-3896**

<u>Google Scholar: Author Only Title Only Author and Title</u>

**Corey E, Matsuda S, Bartel B (1993) Isolation of an Arabidopsis thaliana gene encoding cycloartenol synthase by functional expression in a yeast mutant lacking lanosterol synthase by the use of a chromatographic screen. Proceedings of the National Academy of Sciences 90: 11628-11632**

<u>Google Scholar: Author Only Title Only Author and Title</u>

**Dai L, Liu C, Zhu Y, Zhang J, Men Y, Zeng Y, Sun Y (2015) Functional Characterization of Cucurbitadienol Synthase and**

**Triterpene Glycosyltransferase Involved in Biosynthesis of Mogrosides from Siraitia grosvenorii. Plant Cell Physiol 56: 1172-1182**

<u>Google Scholar: Author Only Title Only Author and Title</u>

**DavidáNes W (1991) Conformational analysis of 10α-cucurbitadienol. Journal of the Chemical Society, Chemical Communications: 1272-1274**

<u>Google Scholar: Author Only Title Only Author and Title</u>

**De La Peña R, Sattely ES (2021) Rerouting plant terpene biosynthesis enables momilactone pathway elucidation. Nature**

**Chemical Biology 17: 205-212**

<u>Google Scholar: Author Only Title Only Author and Title</u>

**Dewick PM (2002) Medicinal natural products: a biosynthetic approach. John Wiley & Sons**

<u>Google Scholar: Author Only Title Only Author and Title</u>

**Dong L, Miettinen K, Goedbloed M, Verstappen FW, Voster A. Jongsma MA, Memelink J, van der Krol S, Bouwmeester HJ (2013) Characterization of two geraniol synthases from Valeriana officinalis and Lippia dulcis: similar activity but difference in subcellular localization. Metabolic engineering 20: 198-211**

<u>Google Scholar: Author Only Title Only Author and Title</u>

**Dong L, Pollier J, Bassard JE, Ntallas G, Almeida A, Lazaridi E, Khakimov B, Arendt P, de Oliveira LS, Lota F, Goossens A Michoux F, Bak S (2018) Co-expression of squalene epoxidases with triterpene cyclases boosts production of triterpenoids in plants and yeast. Metab Eng 49: 1-12**

<u>Google Scholar: Author Only Title Only Author and Title</u>

**Hayashi H, Huang P, Inoue K, Hiraoka N, Ikeshiro Y, Yazaki K, Tanaka S, Kushiro T, Shibuya M, Ebizuka Y (2001) Molecular cloning and characterization of isomultiflorenol synthase, a new triterpene synthase from Luffa cylindrica, involved in biosynthesis of bryonolic acid. European Journal of Biochemistry 268: 6311-6317**

<u>Google Scholar: Author Only Title Only Author and Title</u>

**Hodgson H, De La Pena R, Stephenson MJ, Thimmappa R, Vincent JL, Sattely ES, Osbourn A (2019) Identification of key enzymes responsible for protolimonoid biosynthesis in plants: Opening the door to azadirachtin production. Proc Natl Acad Sci U S A 116: 17096-17104**

<u>Google Scholar: Author Only Title Only Author and Title</u>

**Hussain H, Green IR, Saleem M, Khattak KF, Irshad M, Ali M (2019) Cucurbitacins as Anticancer Agents: A Patent Review. Recent Pat Anticancer Drug Discov 14: 133-143**

<u>Google Scholar: Author Only Title Only Author and Title</u>

**Irmisch S, Jo S, Roach CR, Jancsik S, Man Saint Yuen M, Madilao LL, O’Neil-Johnson M, Williams R, Withers SG, Bohlmann J (2018) Discovery of UDP-Glycosyltransferases and BAHD-Acyltransferases Involved in the Biosynthesis of the Antidiabetic Plant Metabolite Montbretin A. Plant Cell 30: 1864-1886**

<u>Google Scholar: Author Only Title Only Author and Title</u>

**Isaev M (1995) Isoprenoids ofBryonia I. Pentacyclic triterpenes and sterol ofBryonia melanocarpa. Chemistry of Natural Compounds 31: 336-341**

<u>Google Scholar: Author Only Title Only Author and Title</u>

**Itkin M, Davidovich-Rikanati R, Cohen S, Portnoy V, Doron-Faigenboim A, Oren E, Freilich S, Tzuri G, Baranes N, Shen S, Petreikov M, Sertchook R, Ben-Dor S, Gottlieb H, Hernandez A, Nelson DR, Paris HS, Tadmor Y, Burger Y, Lewinsohn E, Katzir**

**N, Schaffer A (2016) The biosynthetic pathway of the nonsugar, high-intensity sweetener mogroside V from Siraitia grosvenorii. Proc Natl Acad Sci U S A 113: E7619-E7628**

<u>Google Scholar: Author Only Title Only Author and Title</u>

**Ito R, Masukawa Y, Hoshino T (2013) Purification, kinetics, inhibitors and CD for recombinant β-amyrin synthase from E uphorbia tirucalli L and functional analysis of the DCTA motif, which is highly conserved among oxidosqualene cyclases. The FEBS journal 280: 1267-1280**

<u>Google Scholar: Author Only Title Only Author and Title</u>

**Kushiro T, Shibuya M, Ebizuka Y (1998) β-Amyrin synthase: cloning of oxidosqualene cyclase that catalyzes the formation of the most popular triterpene among higher plants. European Journal of Biochemistry 256: 238-244**

<u>Google Scholar: Author Only Title Only Author and Title</u>

**Laden BP, Tang Y, Porter TD (2000) Cloning, heterologous expression, and enzymological characterization of human squalene monooxygenase. Archives of biochemistry and biophysics 374: 381-388**

<u>Google Scholar: Author Only Title Only Author and Title</u>

**Li J, Mutanda I, Wang K, Yang L, Wang J, Wang Y (2019) Chloroplastic metabolic engineering coupled with isoprenoid pool enhancement for committed taxanes biosynthesis in Nicotiana benthamiana. Nat Commun 10: 4850**

<u>Google Scholar: Author Only Title Only Author and Title</u>

**Livak KJ, Schmittgen TD (2001) Analysis of relative gene expression data using real-time quantitative PCR and the 2-ΔΔCT method. methods 25: 402-408**

<u>Google Scholar: Author Only Title Only Author and Title</u>

**Martí M, Diretto G, Aragonés V, Frusciante S, Ahrazem O, Gómez-Gómez L, Daròs J-A (2020) Efficient production of saffron crocins and picrocrocin in Nicotiana benthamiana using a virus-driven system. Metabolic Engineering 61: 238-250**

<u>Google Scholar: Author Only Title Only Author and Title</u>

**Nett RS, Lau W, Sattely ES (2020) Discovery and engineering of colchicine alkaloid biosynthesis. Nature 584: 148-153**

<u>Google Scholar: Author Only Title Only Author and Title</u>

**Nett RS, Sattely ES (2021) Total Biosynthesis of the Tubulin-Binding Alkaloid Colchicine. J Am Chem Soc 143: 19454-19465**

<u>Google Scholar: Author Only Title Only Author and Title</u>

**Nguyen L-T, Schmidt HA, Von Haeseler A. Minh BQ (2015) IQ-TREE: a fast and effective stochastic algorithm for estimating maximum-likelihood phylogenies. Molecular biology and evolution 32: 268-274**

<u>Google Scholar: Author Only Title Only Author and Title</u>

**Ohyama K, Suzuki M, Kikuchi J, Saito K, Muranaka T (2009) Dual biosynthetic pathways to phytosterol via cycloartenol and lanosterol in Arabidopsis. Proceedings of the National Academy of Sciences 106: 725-730**

<u>Google Scholar: Author Only Title Only Author and Title</u>

**Otasek D, Morris JH, Bouças J, Pico AR, Demchak B (2019) Cytoscape automation: empowering workflow-based network analysis. Genome biology 20: 1-15**

<u>Google Scholar: Author Only Title Only Author and Title</u>

**Phillips DR, Rasbery JM, Bartel B, Matsuda SP (2006) Biosynthetic diversity in plant triterpene cyclization. Current opinion in plant biology 9: 305-314**

<u>Google Scholar: Author Only Title Only Author and Title</u>

**Rasbery JM, Shan H, LeClair RJ, Norman M, Matsuda SP, Bartel B (2007) Arabidopsis thaliana squalene epoxidase 1 is essential for root and seed development. Journal of Biological Chemistry 282: 17002-17013**

<u>Google Scholar: Author Only Title Only Author and Title</u>

**Reed J, Osbourn A (2018) Engineering terpenoid production through transient expression in Nicotiana benthamiana. Plant Cell Rep 37: 1431-1441**

<u>Google Scholar: Author Only Title Only Author and Title</u>

**Reed J, Stephenson MJ, Miettinen K, Brouwer B, Leveau A, Brett P, Goss RJM, Goossens A, O’Connell MA, Osbourn A (2017) A translational synthetic biology platform for rapid access to gram-scale quantities of novel drug-like molecules. Metab Eng 42: 185-193**

<u>Google Scholar: Author Only Title Only Author and Title</u>

**Ren Y, Kinghorn AD (2019) Natural Product Triterpenoids and Their Semi-Synthetic Derivatives with Potential Anticancer Activity. Planta Med 85: 802-814**

<u>Google Scholar: Author Only Title Only Author and Title</u>

**Sawai S, Saito K (2011) Triterpenoid biosynthesis and engineering in plants. Frontiers in plant science 2: 25**

<u>Google Scholar: Author Only Title Only Author and Title</u>

**Shang Y, Ma Y, Zhou Y, Zhang H, Duan L, Chen H, Zeng J, Zhou Q, Wang S, Gu W, Liu M, Ren J, Gu X, Zhang S, Wang Y, Yasukawa K, Bouwmeester HJ, Qi X, Zhang Z, Lucas WJ, Huang S (2014) Plant science. Biosynthesis, regulation, and domestication of bitterness in cucumber. Science 346: 1084-1088**

<u>Google Scholar: Author Only Title Only Author and Title</u>

**Song W, Yan S, Li Y, Feng S, Zhang J-j, Li J-r (2019) Functional characterization of squalene epoxidase and NADPH-cytochrome P450 reductase in Dioscorea zingiberensis. Biochemical and biophysical research communications 509: 822-827**

<u>Google Scholar: Author Only Title Only Author and Title</u>

**Srisawat P, Fukushima EO, Yasumoto S, Robertlee J, Suzuki H, Seki H, Muranaka T (2019) Identification of oxidosqualene cyclases from the medicinal legume tree Bauhinia forficata: A step toward discovering preponderant α-amyrin-producing activity. New Phytologist 224: 352-366**

<u>Google Scholar: Author Only Title Only Author and Title</u>

**Takase S, Kera K, Hirao Y, Hosouchi T, Kotake Y, Nagashima Y, Mannen K, Suzuki H, Kushiro T (2019) Identification of triterpene biosynthetic genes from Momordica charantia using RNA-seq analysis. Biosci Biotechnol Biochem 83: 251-261**

<u>Google Scholar: Author Only Title Only Author and Title</u>

**Takase S, Kera K, Nagashima Y, Mannen K, Hosouchi T, Shinpo S, Kawashima M, Kotake Y, Yamada H, Saga Y, Otaka J, Araya H, Kotera M, Suzuki H, Kushiro T (2019) Allylic hydroxylation of triterpenoids by a plant cytochrome P450 triggers key chemical transformations that produce a variety of bitter compounds. J Biol Chem 294: 18662-18673**

<u>Google Scholar: Author Only Title Only Author and Title</u>

**Thimmappa R, Geisler K, Louveau T, O’Maille P, Osbourn A (2014) Triterpene biosynthesis in plants. Annu Rev Plant Biol 65: 225-257**

<u>Google Scholar: Author Only Title Only Author and Title</u>

**Wang P, Wei W, Ye W, Li X, Zhao W, Yang C, Li C, Yan X, Zhou Z (2019) Synthesizing ginsenoside Rh2 in Saccharomyces cerevisiae cell factory at high-efficiency. Cell Discov 5: 5**

<u>Google Scholar: Author Only Title Only Author and Title</u>

**Wang P, Wei Y, Fan Y, Liu Q, Wei W, Yang C, Zhang L, Zhao G, Yue J, Yan X, Zhou Z (2015) Production of bioactive ginsenosides Rh2 and Rg3 by metabolically engineered yeasts. Metab Eng 29: 97-105**

<u>Google Scholar: Author Only Title Only Author and Title</u>

**Wang W, Yang H, Li Y, Zheng Z, Liu Y, Wang H, Mu Y, Yao Q (2018) Identification of 16,25-O-diacetyl-cucurbitane F and 25-O-acetyl-23,24-dihydrocucurbitacin F as novel anti-cancer chemicals. R Soc Open Sci 5: 180723**

<u>Google Scholar: Author Only Title Only Author and Title</u>

**Wu S, Jiang Z, Kempinski C, Eric Nybo S, Husodo S, Williams R, Chappell J (2012) Engineering triterpene metabolism in tobacco. Planta 236: 867-877**

<u>Google Scholar: Author Only Title Only Author and Title</u>

**Xu R, Fazio GC, Matsuda SP (2004) On the origins of triterpenoid skeletal diversity. Phytochemistry 65: 261-291**

<u>Google Scholar: Author Only Title Only Author and Title</u>

**Xue Z, Tan Z, Huang A, Zhou Y, Sun J, Wang X, Thimmappa RB, Stephenson MJ, Osbourn A, Qi X (2018) Identification of key amino acid residues determining product specificity of 2,3-oxidosqualene cyclase in Oryza species. New Phytol 218: 1076-1088**

<u>Google Scholar: Author Only Title Only Author and Title</u>

**Yoshida K, Hirose Y, Imai Y, Kondo T (1989) Conformational analysis of cycloartenol, 24-methylenecycloartanol and their derivatives. Agricultural and biological chemistry 53: 1901-1912**

<u>Google Scholar: Author Only Title Only Author and Title</u>

**Zhang Y, Zeng Y, An Z, Lian D, Xiao H, Wang R, Zhang R, Zhai F, Liu H (2022) Comparative transcriptome analysis and identification of candidate genes involved in cucurbitacin IIa biosynthesis in Hemsleya macrosperma. Plant Physiol Biochem 185: 314-324**

<u>Google Scholar: Author Only Title Only Author and Title</u>

**Zhou J, Hu T, Gao L, Su P, Zhang Y, Zhao Y, Chen S, Tu L, Song Y, Wang X (2019) Friedelane-type triterpene cyclase in celastrol biosynthesis from Tripterygium wilfordii and its application for triterpenes biosynthesis in yeast. New Phytologist 223: 722-735**

<u>Google Scholar: Author Only Title Only Author and Title</u>

**Zhou Y, Ma Y, Zeng J, Duan L, Xue X, Wang H, Lin T, Liu Z, Zeng K, Zhong Y, Zhang S, Hu Q, Liu M, Zhang H, Reed J, Moses T, Liu X, Huang P, Qing Z, Liu X, Tu P, Kuang H, Zhang Z, Osbourn A, Ro DK, Shang Y, Huang S (2016) Convergence and divergence of bitterness biosynthesis and regulation in Cucurbitaceae. Nat Plants 2: 16183**

<u>Google Scholar: Author Only Title Only Author and Title</u>

## References

Boykin C, Zhang G, Chen YH, Zhang RW, Fan XE, Yang WM, Lu Q (2011) Cucurbitacin IIa: a novel class of anti-cancer drug inducing non-reversible actin aggregation and inhibiting survivin independent of JAK2/STAT3 phosphorylation. Br J Cancer 104: 781–789

Chen JC, Chiu MH, Nie RL, Cordell GA, Qiu SX (2005) Cucurbitacins and cucurbitane glycosides: structures and biological activities. Nat Prod Rep 22: 386–399

Cho HJ, Tanaka S, Fukui H, Tabata M (1992) Formation of bryonolic acid in cucurbitaceous plants and their cell cultures. Phytochemistry 31: 3893–3896

Corey E, Matsuda S, Bartel B (1993) Isolation of an Arabidopsis thaliana gene encoding cycloartenol synthase by functional expression in a yeast mutant lacking lanosterol synthase by the use of a chromatographic screen. Proceedings of the National Academy of Sciences 90: 11628–11632

Dai L, Liu C, Zhu Y, Zhang J, Men Y, Zeng Y, Sun Y (2015) Functional Characterization of Cucurbitadienol Synthase and Triterpene Glycosyltransferase Involved in Biosynthesis of Mogrosides from Siraitia grosvenorii. Plant Cell Physiol 56: 1172–1182

DavidáNes W (1991) Conformational analysis of 10α-cucurbitadienol. Journal of the Chemical Society, Chemical Communications: 1272–1274

De La Peña R, Sattely ES (2021) Rerouting plant terpene biosynthesis enables momilactone pathway elucidation. Nature Chemical Biology 17: 205–212

Dewick PM (2002) Medicinal natural products: a biosynthetic approach. John Wiley & Sons

Dong L, Miettinen K, Goedbloed M, Verstappen FW, Voster A, Jongsma MA, Memelink J, van der Krol S, Bouwmeester HJ (2013) Characterization of two geraniol synthases from Valeriana officinalis and Lippia dulcis: similar activity but difference in subcellular localization. Metabolic engineering 20: 198–211

Dong L, Pollier J, Bassard JE, Ntallas G, Almeida A, Lazaridi E, Khakimov B, Arendt P, de Oliveira LS, Lota F, Goossens A, Michoux F, Bak S (2018) Co-expression of squalene epoxidases with triterpene cyclases boosts production of triterpenoids in plants and yeast. Metab Eng 49: 1–12

Hayashi H, Huang P, Inoue K, Hiraoka N, Ikeshiro Y, Yazaki K, Tanaka S, Kushiro T, Shibuya M, Ebizuka Y (2001) Molecular cloning and characterization of isomultiflorenol synthase, a new triterpene synthase from Luffa cylindrica, involved in biosynthesis of bryonolic acid. European Journal of Biochemistry 268: 6311–6317

Hodgson H, De La Pena R, Stephenson MJ, Thimmappa R, Vincent JL, Sattely ES, Osbourn A (2019) Identification of key enzymes responsible for protolimonoid biosynthesis in plants: Opening the door to azadirachtin production. Proc Natl Acad Sci U S A 116: 17096–17104

Hussain H, Green IR, Saleem M, Khattak KF, Irshad M, Ali M (2019) Cucurbitacins as Anticancer Agents: A Patent Review. Recent Pat Anticancer Drug Discov 14: 133–143

Irmisch S, Jo S, Roach CR, Jancsik S, Man Saint Yuen M, Madilao LL, O’Neil-Johnson M, Williams R, Withers SG, Bohlmann J (2018) Discovery of UDP-Glycosyltransferases and BAHD-Acyltransferases Involved in the Biosynthesis of the Antidiabetic Plant Metabolite Montbretin A. Plant Cell 30: 1864–1886

Isaev M (1995) Isoprenoids ofBryonia I. Pentacyclic triterpenes and sterol ofBryonia melanocarpa. Chemistry of Natural Compounds 31: 336–341

Itkin M, Davidovich-Rikanati R, Cohen S, Portnoy V, Doron-Faigenboim A, Oren E, Freilich S, Tzuri G, Baranes N, Shen S, Petreikov M, Sertchook R, Ben-Dor S, Gottlieb H, Hernandez A, Nelson DR, Paris HS, Tadmor Y, Burger Y, Lewinsohn E, Katzir N, Schaffer A (2016) The biosynthetic pathway of the nonsugar, high-intensity sweetener mogroside V from Siraitia grosvenorii. Proc Natl Acad Sci U S A 113: E7619–E7628

Ito R, Masukawa Y, Hoshino T (2013) Purification, kinetics, inhibitors and CD for recombinant β-amyrin synthase from E uphorbia tirucalli L and functional analysis of the DCTA motif, which is highly conserved among oxidosqualene cyclases. The FEBS journal 280: 1267–1280

Kushiro T, Shibuya M, Ebizuka Y (1998) β-Amyrin synthase: cloning of oxidosqualene cyclase that catalyzes the formation of the most popular triterpene among higher plants. European Journal of Biochemistry 256: 238–244

Laden BP, Tang Y, Porter TD (2000) Cloning, heterologous expression, and enzymological characterization of human squalene monooxygenase. Archives of biochemistry and biophysics 374: 381–388

Li J, Mutanda I, Wang K, Yang L, Wang J, Wang Y (2019) Chloroplastic metabolic engineering coupled with isoprenoid pool enhancement for committed taxanes biosynthesis in Nicotiana benthamiana. Nat Commun 10: 4850

Livak KJ, Schmittgen TD (2001) Analysis of relative gene expression data using real-time quantitative PCR and the 2-ΔΔCT method. methods 25: 402–408

Martí M, Diretto G, Aragonés V, Frusciante S, Ahrazem O, Gómez-Gómez L, Daròs J-A (2020) Efficient production of saffron crocins and picrocrocin in Nicotiana benthamiana using a virus-driven system. Metabolic Engineering 61: 238–250

Nett RS, Lau W, Sattely ES (2020) Discovery and engineering of colchicine alkaloid biosynthesis. Nature 584: 148–153

Nett RS, Sattely ES (2021) Total Biosynthesis of the Tubulin-Binding Alkaloid Colchicine. J Am Chem Soc 143: 19454–19465

Nguyen L-T, Schmidt HA, Von Haeseler A, Minh BQ (2015) IQ-TREE: a fast and effective stochastic algorithm for estimating maximum-likelihood phylogenies. Molecular biology and evolution 32: 268–274

Ohyama K, Suzuki M, Kikuchi J, Saito K, Muranaka T (2009) Dual biosynthetic pathways to phytosterol via cycloartenol and lanosterol in Arabidopsis. Proceedings of the National Academy of Sciences 106: 725–730

Otasek D, Morris JH, Bouças J, Pico AR, Demchak B (2019) Cytoscape automation: empowering workflow-based network analysis. Genome biology 20: 1–15

Phillips DR, Rasbery JM, Bartel B, Matsuda SP (2006) Biosynthetic diversity in plant triterpene cyclization. Current opinion in plant biology 9: 305–314

Rasbery JM, Shan H, LeClair RJ, Norman M, Matsuda SP, Bartel B (2007) Arabidopsis thaliana squalene epoxidase 1 is essential for root and seed development. Journal of Biological Chemistry 282: 17002–17013

Reed J, Osbourn A (2018) Engineering terpenoid production through transient expression in Nicotiana benthamiana. Plant Cell Rep 37: 1431–1441

Reed J, Stephenson MJ, Miettinen K, Brouwer B, Leveau A, Brett P, Goss RJM, Goossens A, O’Connell MA, Osbourn A (2017) A translational synthetic biology platform for rapid access to gram-scale quantities of novel drug-like molecules. Metab Eng 42: 185–193

Ren Y, Kinghorn AD (2019) Natural Product Triterpenoids and Their Semi-Synthetic Derivatives with Potential Anticancer Activity. Planta Med 85: 802–814

Sawai S, Saito K (2011) Triterpenoid biosynthesis and engineering in plants. Frontiers in plant science 2: 25

Shang Y, Ma Y, Zhou Y, Zhang H, Duan L, Chen H, Zeng J, Zhou Q, Wang S, Gu W, Liu M, Ren J, Gu X, Zhang S, Wang Y, Yasukawa K, Bouwmeester HJ, Qi X, Zhang Z, Lucas WJ, Huang S (2014) Plant science. Biosynthesis, regulation, and domestication of bitterness in cucumber. Science 346: 1084–1088

Song W, Yan S, Li Y, Feng S, Zhang J-j, Li J-r (2019) Functional characterization of squalene epoxidase and NADPH-cytochrome P450 reductase in Dioscorea zingiberensis. Biochemical and biophysical research communications 509: 822–827

Srisawat P, Fukushima EO, Yasumoto S, Robertlee J, Suzuki H, Seki H, Muranaka T (2019) Identification of oxidosqualene cyclases from the medicinal legume tree Bauhinia forficata: A step toward discovering preponderant α-amyrin-producing activity. New Phytologist 224: 352–366

Takase S, Kera K, Hirao Y, Hosouchi T, Kotake Y, Nagashima Y, Mannen K, Suzuki H, Kushiro T (2019) Identification of triterpene biosynthetic genes from Momordica charantia using RNA-seq analysis. Biosci Biotechnol Biochem 83: 251–261

Takase S, Kera K, Nagashima Y, Mannen K, Hosouchi T, Shinpo S, Kawashima M, Kotake Y, Yamada H, Saga Y, Otaka J, Araya H, Kotera M, Suzuki H, Kushiro T (2019) Allylic hydroxylation of triterpenoids by a plant cytochrome P450 triggers key chemical transformations that produce a variety of bitter compounds. J Biol Chem 294: 18662–18673

Thimmappa R, Geisler K, Louveau T, O’Maille P, Osbourn A (2014) Triterpene biosynthesis in plants. Annu Rev Plant Biol 65: 225–257

Wang P, Wei W, Ye W, Li X, Zhao W, Yang C, Li C, Yan X, Zhou Z (2019) Synthesizing ginsenoside Rh2 in Saccharomyces cerevisiae cell factory at high-efficiency. Cell Discov 5: 5

Wang P, Wei Y, Fan Y, Liu Q, Wei W, Yang C, Zhang L, Zhao G, Yue J, Yan X, Zhou Z (2015) Production of bioactive ginsenosides Rh2 and Rg3 by metabolically engineered yeasts. Metab Eng 29: 97–105

Wang W, Yang H, Li Y, Zheng Z, Liu Y, Wang H, Mu Y, Yao Q (2018) Identification of 16,25-O-diacetyl-cucurbitane F and 25-O-acetyl-23,24-dihydrocucurbitacin F as novel anti-cancer chemicals. R Soc Open Sci 5: 180723

Wu S, Jiang Z, Kempinski C, Eric Nybo S, Husodo S, Williams R, Chappell J (2012) Engineering triterpene metabolism in tobacco. Planta 236: 867–877

Xu R, Fazio GC, Matsuda SP (2004) On the origins of triterpenoid skeletal diversity. Phytochemistry 65: 261–291

Xue Z, Tan Z, Huang A, Zhou Y, Sun J, Wang X, Thimmappa RB, Stephenson MJ, Osbourn A, Qi X (2018) Identification of key amino acid residues determining product specificity of 2,3-oxidosqualene cyclase in Oryza species. New Phytol 218: 1076–1088

Yoshida K, Hirose Y, Imai Y, Kondo T (1989) Conformational analysis of cycloartenol, 24-methylenecycloartanol and their derivatives. Agricultural and biological chemistry 53: 1901–1912

Zhang Y, Zeng Y, An Z, Lian D, Xiao H, Wang R, Zhang R, Zhai F, Liu H (2022) Comparative transcriptome analysis and identification of candidate genes involved in cucurbitacin IIa biosynthesis in Hemsleya macrosperma. Plant Physiol Biochem 185: 314–324

Zhou J, Hu T, Gao L, Su P, Zhang Y, Zhao Y, Chen S, Tu L, Song Y, Wang X (2019) Friedelane-type triterpene cyclase in celastrol biosynthesis from Tripterygium wilfordii and its application for triterpenes biosynthesis in yeast. New Phytologist 223: 722–735

Zhou Y, Ma Y, Zeng J, Duan L, Xue X, Wang H, Lin T, Liu Z, Zeng K, Zhong Y, Zhang S, Hu Q, Liu M, Zhang H, Reed J, Moses T, Liu X, Huang P, Qing Z, Liu X, Tu P, Kuang H, Zhang Z, Osbourn A, Ro DK, Shang Y, Huang S (2016) Convergence and divergence of bitterness biosynthesis and regulation in Cucurbitaceae. Nat Plants 2: 16183

